# Non-overlapping social and food reward representations in the basolateral amygdala

**DOI:** 10.1101/2025.10.06.680770

**Authors:** Jarildy L. Javier, Hymavathy Balasubramanian, Jennifer Isaac, Larry J. Young, Malavika Murugan

**Affiliations:** Emory Neuroscience Graduate Program, Emory University, Atlanta, GA 30322, USA; Center for Translational Social Neuroscience, Emory University, Atlanta, GA 30329, USA; Silvio O. Conte Center for Oxytocin and Social Cognition, Emory National Primate Research Center, Emory University, Atlanta, GA 30329, USA; Department of Biology, Emory University, Atlanta, GA 30322, USA

**Author notes:** **Corresponding author:** Malavika Murugan**, email:**.

## Abstract

The ability to consider and appraise positively valenced stimuli in the environment, such as food and social interaction, to guide appropriate action is important for survival of most animals. Several studies have compared how food and social rewards are represented in different regions involved in reward processing and found either overlapping or distinct representations. In the basolateral amygdala (BLA) there seems to be opposing evidence for both shared and unique encoding of social and nonsocial stimuli. In our recent work, we found that the medial prefrontal cortex (mPFC), a region reciprocally connected to the BLA, has distinct social and food representations using a novel self-paced two-choice assay. Given the BLA and mPFC’s differing roles in reward processing, it is important to understand how these two nodes may differ in their encoding reward types within the same assay. To resolve how the BLA encodes social and food information, we recorded the activity of individual BLA neurons in female and male mice during a two-choice social-sucrose operant task. We found that BLA neurons robustly and distinctly respond to social and food reward. In contrast to the mPFC, BLA neurons did not show a bias towards social reward responsiveness and instead showed equal social/sucrose representation, in males, or a sucrose reward bias, in females. BLA neurons were sensitive to internal state - water deprivation increased the proportion of sucrose reward responsive neurons. Additionally, sucrose reward responsive BLA neurons were differentially sensitive to reward omissions, such that neurons that were excited by sucrose reward were more sensitive to reward omissions compared to those inhibited by reward. Together, these findings demonstrate distinct, heterogeneous response profiles within the BLA to social and food rewards, in a manner different from the mPFC.

## Introduction

A wide range of naturally occurring stimuli, such as food and conspecifics, are positively reinforcing and can elicit robust reward-seeking behaviors in animals. Several factors, both external (e.g., resource availability, proximity) and internal (e.g., thirst, isolation), strongly influence decisions to pursue one reward over another. Deciding how to allocate time and effort among these options requires the brain to encode the value of different stimuli, compare them, and guide adaptive choice. Disruptions in this process are thought to contribute to neuropsychiatric disorders, including depression^1–7^, addiction^8–15^, and schizophrenia^16–22^, making it essential to understand how reward information is represented across modalities in the brain.

There has been a lot of recent interest in understanding how social and food rewards are represented in various nodes of the reward circuitry^23^. Studies across species indicate that social and nonsocial rewards recruit overlapping mesolimbic networks^24–31^. Although it is unclear whether these reward types are represented by the same neurons or by distinct neuronal populations within reward-related regions. For example, it appears that dopamine neurons in the ventral tegmental area (VTA), a key node in supporting reward-related behaviors, found overlapping populations of neurons to respond to social and food stimuli^31^. In contrast, in recent work, we found that largely non-overlapping populations of medial prefrontal cortex (mPFC) neurons are responsive to social and food stimuli in mice freely choosing between social and food rewards in a two-choice social-sucrose operant assay^25,32^. However, there is debate on how social and non-social information is represented in the basolateral amygdala (BLA), a region with strong connections with the mesolimbic reward circuit.

The BLA has been heavily implicated in valence encoding and in processing saliency information^33–41^. The BLA is strongly recruited by both positive and negative valence stimuli, specifically non-social stimuli^36,42–52^. Work has also shown the BLA to be modulated during both positive^53,54^ and negative social interactions^55–61^. While we have learned much about how the BLA encodes positive versus negative valence stimulus^34,39,42–45,47,62–71^, we still know relatively little about how two stimuli with positive valence are represented in the BLA. For example, distinct BLA sub-populations represent stimulus-specific value of two different food rewards^72^. It is unknown if these findings would extend to social and non-social rewards. In other words, it remains unclear whether the BLA uses the same neural populations^44,73,74^ or distinct populations^23,75–77^ to represent rewards of different types.

Evidence supports both possibilities. In head-fixed macaques, decoders trained on amygdala responses to juice magnitude could also classify social hierarchy, suggesting that amygdala neurons multiplex social and nonsocial information^75,78^. In this case, however, the social stimuli (faces of other monkeys) served as conditioned cues, whereas juice rewards of varying volumes acted as the unconditioned reward. In contrast, in rats, BLA populations responding to social or food stimuli were largely distinct when rats were allowed to interact with rewards present in an arena^79^. However, it is hard to dissociate the appetitive and consummatory components of social behavior during passive, free-range interactions. Given BLA’s role in modulating a range of social behaviors, it would be beneficial to record the activity of BLA neurons when animals can freely and actively choose between social and non-social rewards.

Additionally, because the mPFC and BLA are heavily reciprocally interconnected nodes in reward-processing networks but are thought to serve distinct functions^23,80,81^, it is important to determine whether the BLA and mPFC show a similar or different coding scheme under identical behavioral conditions. The mPFC is important for encoding reward and social context related information^25,45,46,83–88^, and both the mPFC and BLA send glutamatergic inputs to the NAc, known to drive reward-seeking behaviors^23,89–91^, promoting intracranial self-stimulation^89,92–98^. Projections between the mPFC and BLA have been shown to regulate social novelty preference^99–102^, social avoidance^103,104^, social hierarchy^78,105–107^, food related behaviors ^108–113^, and behavioral flexibility to reward and aversive stimuli^110,114–118^. Thus, it is clear that both regions are critically involved in social and nonsocial reward information processing - but comparing how exactly these two regions process reward can pose challenges when comparing across paradigms. In this study, we investigate the role of the BLA in social and food reward processing and leverage the two-choice social-sucrose assay to directly compare reward encoding across these two nodes.

Here, we used in vivo calcium imaging to record the activity of individual BLA neurons in freely moving male and female mice performing the two-choice social–sucrose operant assay. We found that largely non-overlapping populations of BLA neurons represent social and non-social choice and reward information. We discovered that BLA neurons are less modulated by social information when compared to mPFC neurons under similar experimental conditions. We found that BLA neurons are sensitive to changes in internal state, with a much larger fraction of neurons responding to the sucrose-water reward in water-restricted mice compared to mice that had *ad libitum* access to water. Finally, we found that BLA neurons that are excited in response to sucrose reward are more sensitive to sucrose reward omissions in comparison to those that are inhibited in response to sucrose reward. Taken together, our findings show that social and non-social rewards recruit non-overlapping populations of BLA neurons. Despite being strongly reciprocally connected, the BLA and mPFC exhibit differences in how they represent social and non-social reward information.

## Results

### BLA neurons robustly respond to both social and food rewards

To directly compare social and nonsocial food reward representations, we first trained female and male mice on a self-paced two-choice operant assay which we recently developed^25,32^. Mice were allowed to freely choose between access to a novel, same-sex conspecific or a sucrose reward (10 µL drop of 10% sucrose in water solution) (Figure 1A, n=10 female mice, 7 male mice).

**Figure 1:**
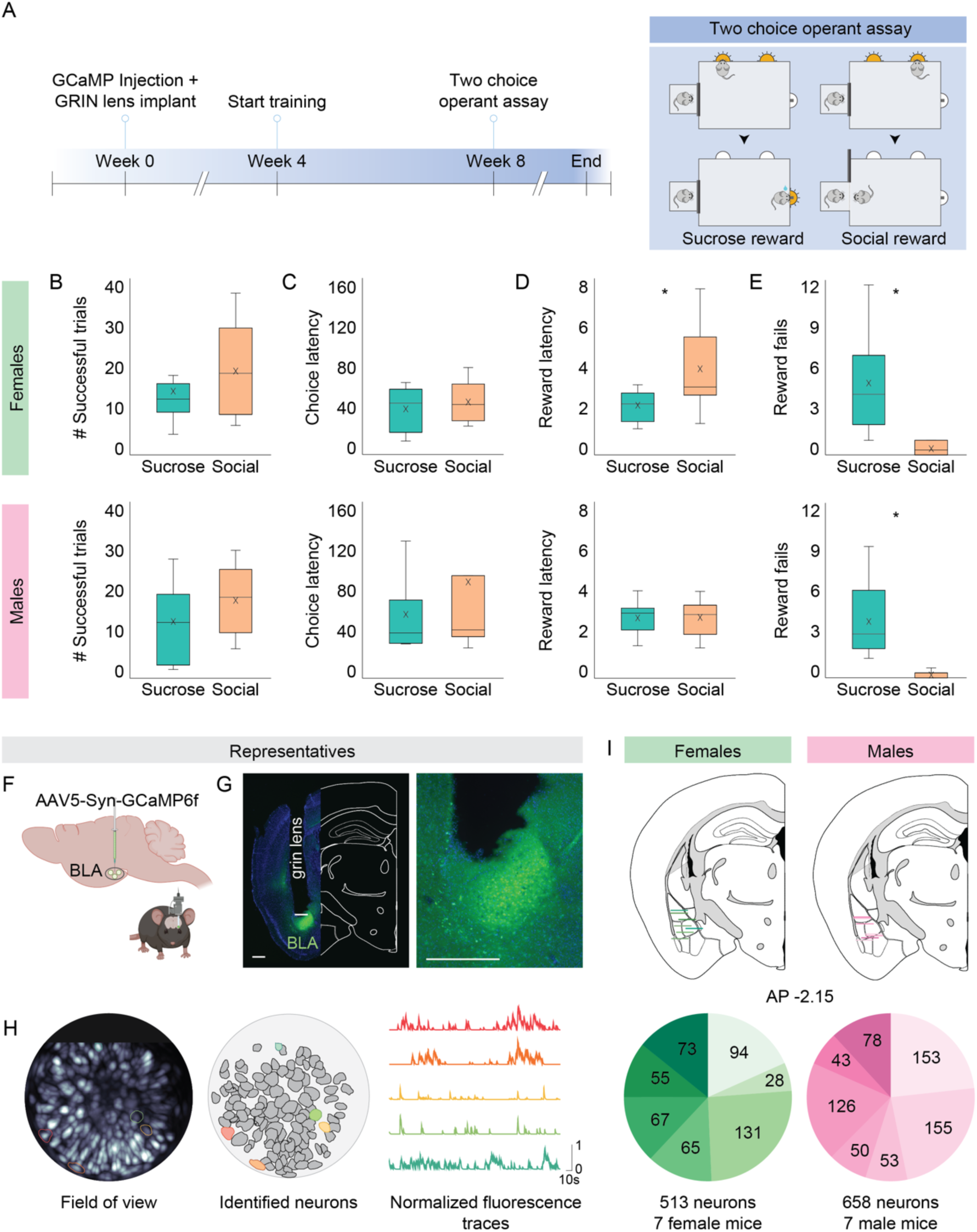
Imaging basolateral amygdala neurons while mice choose between food and social rewards. A. Experimental timeline (left) and schematic (right) of the two-choice operant assay. Fully trained mice can freely choose between a sucrose or social choice port to obtain either 10 μL of 10% sucrose solution or access to a novel same-sex conspecific for 20 seconds, respectively. B. Fully trained female (top row) and male mice (bottom row) made an equivalent number of social and sucrose trials (paired t-test, number of trials: female: p=0.410, male: p=0.350). C. Both females and males displayed similar choice latencies for social and sucrose trials (paired t-test, choice latency: female: p=0.370, male: p=0.302). D. While females had a slightly increased social reward latency compared to sucrose, males showed comparable social and sucrose reward latencies (paired t-test, reward latency: female: p=0.020, male: p=0.965). E. Both females and males made more sucrose reward fails compared to social reward fails (paired t-test, reward fails: female: p=0.002, male: p=0.008). N=10 female mice, 7 male mice, 3 behavioral sessions per mouse. For panels B-E top row indicates data from female mice and bottom row indicates data from male mice. Box plot center lines denote the median, the X’s denote the mean, and the box edges denote the 25th and 75th percentiles. The whiskers extend to minimum and maximum values, excluding any outliers that extend 1.5x the interquartile range above the 75th or below the 25th percentile. F. Schematic of the viral strategy. G. Representative histology showing GCaMP expression and GRIN lens placement in the BLA. Scale bar: 500 um. H. Example field of view during cellular resolution calcium imaging of basolateral amygdala neurons during the two-choice operant assay. I. Reconstruction of GRIN lens placement for calcium imaging in the basolateral amygdala for females (left) and males (right). Each colored line corresponds to the same-colored slice in the pie chart and shows the position of the lens at an AP of −2.15 relative to bregma in the Allen Mouse Brain Common Coordinate Framework. We recorded the activity of individual BLA neurons in 7 female (n=513 neurons) and 7 male (n=658 neurons) mice during the two-choice (social-sucrose) operant task. Schematic in A adapted from Isaac & Balasubramanian, et al. (2025). Viral injection brain and mouse schematic in F created with BioRender.com released under a Creative Commons Attribution-NonCommercial-NoDerivs 4.0 International License (https://creativecommons.org/licenses/by-nc-nd/4.0/deed.en.)

After training, both females and males were equally motivated by the social and sucrose rewards, successfully completing similar numbers of social and sucrose trials (Figure 1B, paired t-test, female: p=0.410, male: p=0.350). Mice also showed similar choice latencies for social and sucrose trials (Figure 1C, paired t-test, female: p=0.370, male: p=0.302). Females had slightly longer social reward latencies compared to sucrose, while males had no difference (Figure 1D, paired t-test, female: p=0.020, male: p=0.965). Both females and males made fewer social reward fails than sucrose reward fails (Figure 1E, paired t-test, female: p=0.002, male: p=0.008). These behavioral results are in line with a previous study employing the same two-choice task^25,32^, showing that during full *ad libitum* water access conditions mice will make equivalent numbers of pokes for social and sucrose reward.

To explore how social and nonsocial food reward is represented by the BLA, we recorded the activity of individual BLA neurons in fully trained female and male mice during the two-choice task (Figure 1F-I, n=7 female mice, 7 male mice). We expressed a genetically encoded calcium sensor and imaged the activity of over 1100 neurons BLA neurons across 14 mice via GRIN lens using an *in vivo* one photon miniscope (Figure 1F-I, female n=513 neurons, 7 mice, male n=658 neurons, 7 mice). Recorded calcium activity was aligned to five task events during the two-choice assay: trial start, social choice, sucrose choice, social reward and sucrose reward (Figure 2A). Then, we identified individual BLA neurons that were significantly modulated by each task event (Figure 2B). While males had similar proportions of sucrose and social responsive neurons (Figure 2D, E; proportion z-test, p=0.1471), females had greater sucrose reward representation compared to social reward (Figure 2C, E; proportion z-test, p=0.0005). We also found greater sucrose reward representation compared to social reward in an encoding model based on BLA neural activity for both females and males (Supp. Figure 1D; proportion z-test, social reward vs sucrose reward, female: p<0.00001; male: p=0.0005).

**Figure 2:**
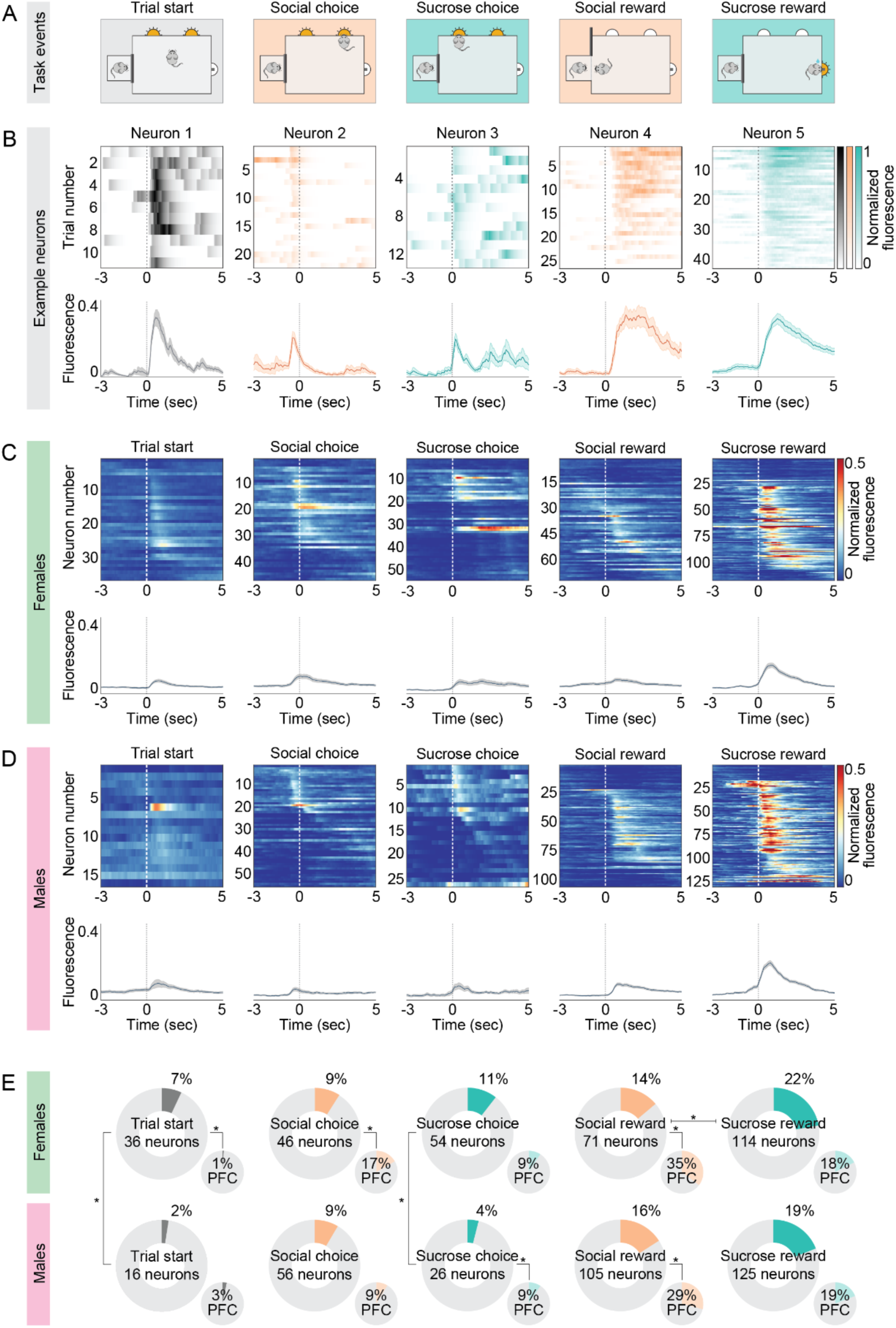
BLA neurons represent sucrose and social reward in similar proportions, in contrast to the strong social reward preference of the mPFC. A. Schematic of two-choice operant assay task events to which BLA neuron activity was aligned, from left to right: trial start, social choice, sucrose choice, social reward, sucrose reward. B. Examples of trial-by-trial responses of individual neurons that were significantly modulated (excited or inhibited) by each of the task events. The top row displays heat maps of normalized trial-by-trial activity for each neuron. Bottom row shows average normalized fluorescence across all trials for each neuron. A dashed line at zero denotes the onset of each task event. The following time windows relative to event onset were considered for determining significance: trial start −1.5s to 1.5s; choice −2.5s to 0.5s; reward −0.5s to 2.5s. C, D. Top row: Heatmaps showing average activity of all significant neurons across trials for various task events. Neurons are sorted by the time of maximum fluorescence across each task event. Neurons that are classified as modulated by more than one task event are included in all relevant heatmaps. Bottom row: The average fluorescence traces of all significantly responsive neurons from the corresponding heatmap. Shaded error regions indicate ± SEM. C displays data from females, and D from males. E. In large donut charts are the proportions of total recorded basolateral amygdala neurons that are modulated by each task event in females and males (female: n=513 neurons, 7 mice; male: n=658 neurons, 7 mice). In the small pie charts are the proportions of mPFC neurons modulated by each task event in the two-choice operant assay from Isaac et al. (2024). Females and males seem to represent task parameters in similar proportions for all except trial start and sucrose choice. A smaller fraction of BLA neurons in male mice respond to trial start and sucrose choice (proportion z-test, female vs male: trial start: p=0.00016, social choice: p=0.787, sucrose choice: p<0.00001, social reward: p=0.313, sucrose reward: p=0.174). Unlike the mPFC, which has a demonstrated social reward representation bias in the two-choice assay, in males BLA neurons seem to similarly represent sucrose and social reward and in females there is a bias toward sucrose reward not social reward (proportion z-test, social reward vs sucrose reward, female: p=0.0005; male: p=0.1471). Indeed, proportionally less BLA neurons responded to social reward compared to the mPFC in females and males, with females also having a lower fraction of social choice neurons compared to the mPFC (proportion z-test, BLA vs mPFC, female: trial start: p<0.00001, social choice: p<0.00001, sucrose choice: p=0.497, social reward: p<0.00001, sucrose reward: p=0.0536; male: trial start: p=0.401, social choice: p=0.904, sucrose choice: p=0.0002, social reward: p<0.00001, sucrose reward: p=0.944). Schematic in A adapted from Isaac et al. (2024). The Bonferroni corrected threshold for proportion comparisons for trial start, social and sucrose choice is p=0.025 (female vs male, BLA vs mPFC), for social and sucrose reward it’s p=0.017 (female vs male, BLA vs mPFC, social vs sucrose).

In order to determine how neural representations compare across another key node in the reward circuit, we compared responses of BLA neurons to responses of mPFC neurons acquired in the same task^25^. Interestingly, we found that there is a much more muted response in BLA to social reward compared to the mPFC, with a smaller fraction of BLA neurons responding to social reward relative to the mPFC (proportion z-test, female: BLA 14% vs mPFC 35%, p<0.00001, male: BLA 16% vs mPFC 29%, p<0.00001). These results suggest that different nodes involved in social behavior and reward processing can differentially encode social and food rewards, even when strongly reciprocally connected, like the BLA and mPFC are in this case^67,81,103^.

### Largely non-overlapping populations of BLA neurons represent social and sucrose reward

BLA neurons are thought to broadly encode valence information^36–40,42,43^, therefore we explored the possibility that the same neurons in the BLA could respond to both social and sucrose reward, both positively reinforcing stimuli. To do so, we directly compared responses of BLA neurons to social and sucrose choice and reward (Figure 3). First, we found that more BLA neurons encoded reward compared to choice in both male and female mice (Figure 3A). In categorizing the reward response further, we found that BLA neurons are more likely to be positively modulated, i.e., increase their activity, during sucrose reward consumption rather than inhibited (Figure 3B, proportion z-test, female: excite: 16%, inhibit: 6%, p<0.00001; male: excite: 16%, inhibit: 3%, p<0.00001). This bias in excitation was not as evident in the social reward response, with females having similar fractions of social reward excite and inhibit neurons and males showing a significant but relatively smaller bias towards social reward excite compared to sucrose reward (Figure 3C, proportion z-test, female: excite: 8%, inhibit: 6%, p=0.1096; male: excite: 10%, inhibit: 6%, p=0.00596). In contrast, in the mPFC, neurons were more likely to show increased activity in response to social reward and rarely inhibited^25^. Furthermore, in contrast to the BLA, a similar proportion of mPFC neurons are excited or inhibited in response to sucrose reward^25^. These findings also suggest that social and food reward information is differentially organized in the BLA and the mPFC.

**Figure 3:**
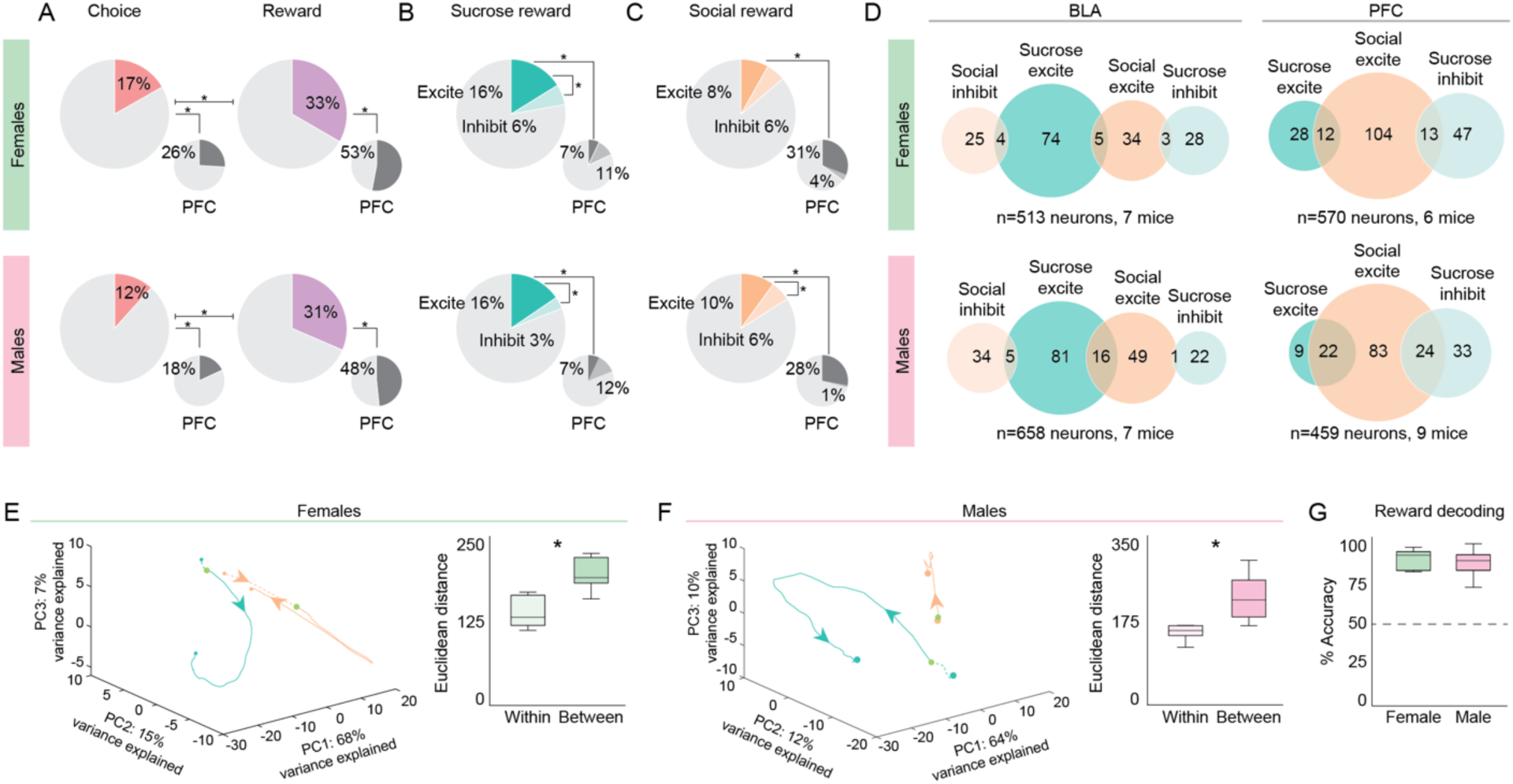
The BLA represents social and sucrose rewards through largely non-overlapping populations of neurons. A. In large pie charts are the proportions of total recorded basolateral amygdala neurons that are significantly responsive to choice (left) and reward (right), generally, sucrose reward and social reward in females (top row; n=513 neurons, 7 mice;) and males (bottom row; n=658 neurons, 7 mice). In the small pie charts are the proportions of mPFC neurons in each respective group from Isaac et al 2024. In the BLA, there are more reward responsive neurons compared to choice for both males and females (proportion z-test, female: choice: 86/513, reward: 171/513, p<0.00001; male: choice: 77/658, reward: 207/658, p<0.00001), a pattern also observed in the mPFC. The Bonferroni corrected threshold for proportion comparisons is p=0.025 (choice vs reward, BLA vs mPFC). B. A higher proportion of BLA neurons are excited (positively modulated) than inhibited by sucrose reward (proportion z-test, female: excite: 83/513, inhibit: 31/513, p<0.00001; male: excite: 102/658, inhibit: 23/658, p<0.00001). In both males and females, there is a higher proportion of sucrose reward excite neurons in the BLA compared to the mPFC (proportion z-test, female: BLA: 83/513, mPFC: 40/570, p<0.00001; male: BLA: 102/658, mPFC: 31/459, p<0.00001). The Bonferroni corrected threshold for proportion comparisons for sucrose reward excite is p=0.025 (excite vs inhibit, BLA vs mPFC), for inhibit it’s p=0.05 (excite vs inhibit). C. In males but not females, there were more social reward excite than social reward inhibit neurons in the BLA (proportion z-test, female: excite: 42/513, inhibit: 29/513, p=0.1096; male: excite: 66/658, inhibit: 39/658, p=0.00596). Additionally, there are proportionately more social reward excite neurons in the mPFC than the BLA (proportion z-test, female: BLA: 42/513, mPFC: 177/570, p<0.00001; male: BLA: 66/658, mPFC: 129/459, p<0.00001). The Bonferroni corrected threshold for proportion comparisons for social reward excite is p=0.025 (excite vs inhibit, BLA vs mPFC), for inhibit it’s p=0.05 (excite vs inhibit). D. Left column: Venn diagrams displaying the overlap of neurons that are either excited or inhibited by sucrose and social reward in the BLA and mPFC. Circle sizes are proportional to the number of neurons in each group. In both male and female mice, neurons that were positively modulated by sucrose reward did so almost exclusively, with few sucrose excite neurons also responding to social reward (proportion z-test, female: sucrose excite only: 74/83 89.16%, sucrose excite and social reward responsive: 9/83 10.84%, p<0.00001; male: sucrose excite only: 81/102 79.41%, sucrose excite and social reward responsive: 21/102 20.38%, p<0.00001). This pattern and group exclusivity is mirrored in the mPFC (right column), but with social reward as the largest class. Not shown: BLA overlap between social inhibit and sucrose inhibit for females is 2 neurons, for males is 1 neuron. E. Left: Principal component analysis of trial-averaged population neural activity for both sucrose (teal) and social (orange) reward trials for females (n=513 neurons). Arrowhead indicates direction of time. The filled green circle indicates reward onset. Right: The Euclidean distance separating the PC-projected population vectors in the reward window is significantly greater between social and sucrose trials than within each trial type (unpaired two sample t-test, within vs between, p=0.0001). Shaded error regions indicate +/− SEM. F. Same as C, trial-averaged population traces for sucrose and social reward trials plotted on the first 3 principal components in state space for males (n=658 neurons). Euclidean distance between social and sucrose trial population vectors was greater than within trial type (unpaired two sample t-test, within vs between, p=0.0021). G. Decoders trained on either female or male neural data were equally able to distinguish between social and sucrose rewards (unpaired two sample t-test, female vs male, p=0.6952; paired two sample t-test, female vs shuffled, p<0.00001, male vs shuffled, p<0.00001). Decoding accuracy was calculated for each animal from all recorded neurons, with a trial-matched number of sucrose and social trials. All decoding was significantly greater than shuffled data, indicated by a dashed line at 50%. Box plot center line indicates the median, box edges indicate the 25th and 75th percentile and whiskers extend to +/− 2.7x the standard deviation. N=7 female mice, 7 male mice.

Given that BLA neurons are known to multiplex social and food reward-related information in primates^78^, we determined if individual neurons are responsive to both social and food rewards during active reward-seeking. We determined the overlap in response groups between reward types. We found that the BLA, like the mPFC, seems to represent reward through mostly non-overlapping groups of neurons (Figure 3D). Indeed, most reward responsive neurons in the BLA seem to be specialized towards a specific reward response (Supp. Figure 3).

Next, to determine whether sucrose and social reward representations in the BLA are separable and decodable at the population level, we plotted the neural population trajectories of social and sucrose reward responses (Figure 3E-F). For both females and males, reward trajectories within a reward type were more similar to each other than across reward types, as determined by comparisons of pairwise Euclidean distances of population vectors within and across trial-types, supporting that differences observed at the individual neuron level also extends to the population level. Additionally, a decoder trained on reward responses was successfully able to assign a neural reward response to the correct reward type with high accuracies and significantly above chance for both females and males (Figure 3E, unpaired two sample t-test, female vs male, p=0.6952; paired two sample t-test, female vs shuffled, p<0.00001, male vs shuffled, p<0.00001). Taken together, these data suggest that sucrose and social representations in the BLA are unique and decodable at both the individual neuron and population level.

Choice responsive neurons were also found to be partly non-overlapping (Supp. Figure 4D) and specialized in their responses (Supp. Figure 2). While comparisons of pairwise Euclidean distances of population vectors within and across trial-types of the choice data demonstrated that the population trajectories of sucrose choice and social choice neurons were separable (Supp. Figure 4E-F), choice decoding was not significantly above chance (Supp. Figure 4G-H) suggesting that at the population-level social and sucrose choice information is weakly separable in the BLA.

### Sucrose reward representations in the BLA are modulated by internal state

Given the large contribution of sucrose reward excite neurons to overall reward representations in the BLA (Figure 3A-D), we next sought to investigate whether sucrose reward representations would be flexible to changes in internal state and reward omissions. To determine if BLA neurons are sensitive to internal state changes, we placed fully trained mice on a restricted water schedule, ran them on the two-choice assay, and assessed changes in behavioral performance and neural activity (Supp. Figure 5A). As expected, restricted water access caused mice to pursue more sucrose rewards than social rewards (Supp. Figure 5B). Additionally, mice chose the sucrose reward more during restricted water access than when they had full *ad libitum* access to water (Supp. Figure 5B, two-factor anova, factors: water condition (FW/RW), trial type (sucrose/social): female: interaction p<0.00001, water condition p<0.00001, trial type p<0.00001; male: interaction p<0.00001, water condition p<0.00001, trial type p<0.00001; post-hoc unpaired t-test comparing FW/RW within trial type: female: sucrose: p<0.00001, social: p=0.0013; male: sucrose: p<0.00001, social: p=0.0466).

Compared to the full water access condition, a greater proportion of BLA neurons responded to sucrose reward during restricted access to water in female and male mice (Supp. Figure 6D, proportion z-test, RW vs FW, sucrose reward, female: p<0.00001, male: p<0.00001). Additionally, proportionately more BLA neurons responded to sucrose reward compared to social reward (Supp. Figure 6D, proportion z-test, social vs sucrose reward, RW: female: p<0.00001, male: p<0.00001, FW: female: p=0.0005; male: p=0.1471). Female mice had more BLA neurons significantly modulated by sucrose reward compared to male mice during restricted water access (Supp. Figure 6D, proportion z-test, female vs male: p<0.00001). These results demonstrate that food reward responsive BLA neurons are sensitive to internal state changes.

### Sucrose reward representations in the BLA are modulated by the absence of reward

Next, to determine if BLA sucrose reward representations are sensitive to the presence of reward once the association between cue, choice and reward has been learned, we placed a subset of mice through a modified two-choice assay where on ∼30% of sucrose reward trials reward was omitted (Supp. Figure 7A, n= 5 mice, 3 female, 2 male). During the sucrose omission two-choice assay, mice displayed typical two-choice behavior (Supp. Figure 7B). Even when sucrose reward was unreliable, BLA neurons still responded robustly to both social and sucrose rewards, with higher representation of sucrose compared to social reward (Supp. Figure 7C-E, proportion z-test, p<0.00001). BLA neurons also more highly represented sucrose reward during the sucrose omission session compared to the full water condition (Supp. Figure 7E, proportion z-test, p<0.00001).

To understand how sucrose reward responsive neurons may respond to the absence of an expected sucrose reward, we compared neuron activity during the reward period between rewarded and unrewarded trials (Figure 4). We found that 63% of sucrose reward excite neurons became significantly less responsive during the absence of reward (Figure 4A-B, 76/122 sucrose excite neurons, paired t-test, rewarded vs unrewarded population response: p<0.00001). While the remaining sucrose reward excite neurons did not individually show significant differences in fluorescence between rewarded and unrewarded trials, these “non-modulated” neurons did show a decrease in activity during the unrewarded trials at the population level (Figure 4C, 46/122 sucrose excite neurons, paired t-test, rewarded vs unrewarded population response: p<0.00001).

**Figure 4:**
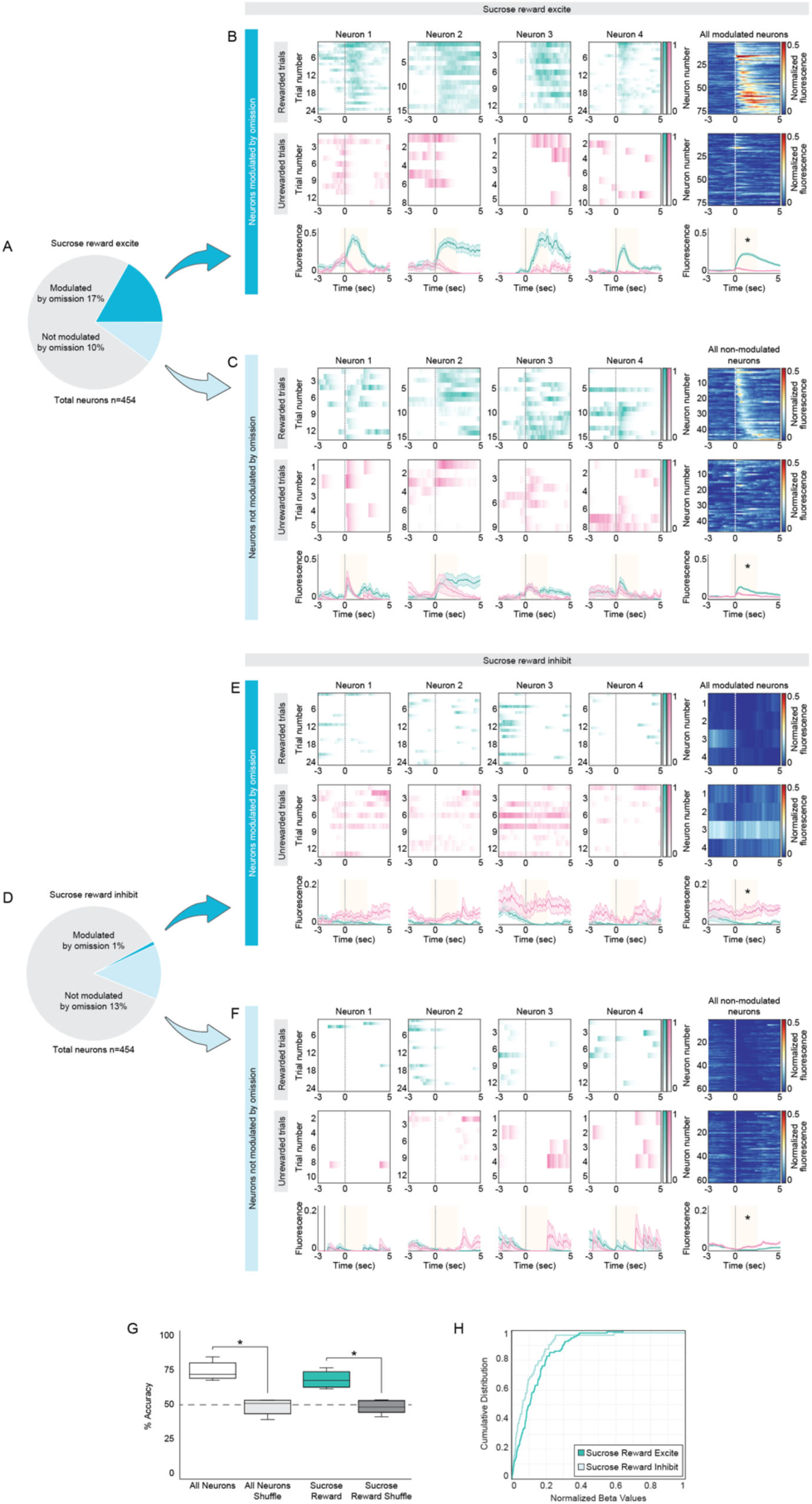
Sucrose reward responsive neurons in the BLA are significantly modulated by the absence of an expected sucrose reward. A. Mice underwent a modified version of the two-choice operant assay (n=5 mice, 3 female, 2 male). On ∼30% of sucrose trials, the sucrose reward was withheld. Neural responses within the reward window were compared between rewarded and unrewarded trials for neurons significantly responsive to sucrose reward (reward window: −0.5s to 2.5s relative to reward consumption onset). Of 454 total neurons, 186 were sucrose reward responsive, 122 of which were excited by sucrose reward, increasing their activity in response to sucrose. During unrewarded trials, 76 neurons (17% of total neurons) significantly decreased their activity compared to rewarded trials while 46 neurons (10%) were not significantly modulated by the absence of reward (two tailed t-test, modulation threshold p=0.05). B. Neuronal data for sucrose reward excite neurons significantly modulated by absence of reward. The first four columns show responses of four example neurons and the rightmost column displays data from all modulated sucrose reward excite neurons (n=76 neurons). Heatmaps display the average normalized fluorescence during rewarded (top row) and unrewarded (middle row) sucrose trials for individual neurons and for the population. Population heatmap for rewarded trials displays neurons sorted by time of maximum fluorescence. Population heatmap for unrewarded trials preserves the same order as the rewarded trials heatmap, such that neurons remain on the same row between both maps. The bottom row shows the average normalized fluorescence traces for rewarded (teal) and unrewarded (pink) trials of each respective set of heatmaps. A dashed line at zero shows sucrose reward consumption onset. Shaded error regions indicate ± SEM. Yellow highlights the reward window. In the absence of reward there is a decrease in activity evident at both the individual neuron (paired t-test, neuron 1-4 p=0.0025, 0.0002, 0.0028, 0.0001) and the population level (paired t-test, p<0.00001). C. Neural data for sucrose reward excite neurons that were not significantly modulated by withheld reward. Though individual neurons did not reveal a significant response to sucrose reward absence (paired t-test, neuron 1-4 p=0.4855, 0.5131, 0.4288, 0.5265), at the population level there emerged a marked decrease in activity following reward withholding (paired t-test, p<0.00001). D. Of 186 sucrose reward responsive neurons, 64 neurons were inhibited by sucrose reward, decreasing their activity in response to sucrose. During unrewarded trials, only 4 neurons (1% of total neurons) significantly increased their activity compared to rewarded trials, and 60 neurons (13%) were not significantly modulated by the absence of reward (two tailed t-test, modulation threshold p=0.05). E. Neuronal data for sucrose reward inhibit neurons significantly modulated by absence of reward (n=4 neurons). Population heatmap for rewarded trials displays neurons sorted by time of minimum fluorescence. Population heatmap for unrewarded trials preserves the same order as the rewarded trials heatmap, such that neurons remain on the same row between both maps. Yellow highlights the reward window. During unrewarded trials (pink), neurons show more elevated activity relative to rewarded trials (teal) (paired t-test: neuron 1-4 p=0.0.0316, 0.0.0032, 0.0085, 0.0065; population p<0.00001). F. Neural data for sucrose reward inhibit neurons that were not significantly modulated by withheld reward. Though individual neurons did not reveal a significant response to sucrose reward absence (paired t-test, neuron 1-4 p=1.000, 0.9596, 1.000, 0.5513), at the population level there was a loss of inhibition during unrewarded trials (paired t-test, p<0.00001). |G. A decoder trained on neural responses from all neurons (n= 5 mice, 454 neurons) was able to successfully distinguish between rewarded and unrewarded sucrose trials (paired two sample t-test, all neurons vs shuffled, p=0.0095) on the test set (70-30 train-test ratio). Similarly, when the decoder was limited to responses from only sucrose reward responsive neurons, the decoder still performed well above chance (n=5 mice, 186 neurons; paired two sample t-test: all neurons vs sucrose reward, p=0.1666; sucrose reward vs shuffled, p=0.0114). Decoding accuracy was calculated per animal with a trial-matched number of unrewarded and rewarded sucrose trials. The dashed line indicates 50% accuracy. The Bonferroni corrected threshold for decoding accuracy comparisons is p=0.025 (actual vs shuffled, all neurons vs sucrose reward). H. Cumulative distribution plot of normalized beta values for sucrose reward excite and inhibit neurons from the sucrose reward neuron-based decoder (teal in G). Normalized betas from sucrose reward excite neurons are overall higher than sucrose reward inhibit neurons, indicated by a curve shift to the right (Kolmogorov-Smirnov test for normality: excite n=122 neurons, p<0.00001, inhibit n=64 neurons, p<0.00001; Mann-Whitney U test: excite vs inhibit, p=0.0002).

For BLA neurons that were typically inhibited in response to sucrose reward, only a very small subset of those neurons was significantly differentially modulated by sucrose reward omission (Figure 4D-E, 4/64 sucrose inhibit neurons, paired t-test, rewarded vs unrewarded population response: p<0.00001). The rest of the sucrose reward inhibited neurons, while not showing effects at the individual neuron level, showed an effect at the population level. These “non-modulated” sucrose reward inhibit neurons showed more elevated activity during the reward period during reward omission trials relative to rewarded trials (Figure 4F, paired t-test, rewarded vs unrewarded population response: p<0.00001). These data support the idea that BLA sucrose reward neurons are extremely sensitive to the presence of the expected reward.

Finally, we examined whether these population response differences to the presence and absence of expected sucrose reward were decodable. We built two different decoders, one was trained on neural responses during rewarded and unrewarded sucrose trials from all neurons, another was restricted to responses from just sucrose reward responsive neurons. Both decoders performed similarly; we were able to correctly decode the identity of rewarded and unrewarded trials with high accuracy and significantly above chance from BLA activity (Figure 4G, n=5 mice, 186 neurons; paired two sample t-test: all neurons vs shuffled, p=0.0095, sucrose reward vs shuffled, p=0.0114; all neurons vs sucrose reward, p=0.1666).

This suggests that activity of the sucrose reward responsive population in the BLA alone is sufficient to convey the presence or absence of reward to downstream regions. To determine how sucrose reward excite and inhibit neurons may be contributing to this decodability, we assessed the beta values of individual sucrose reward excite and inhibit neurons from the sucrose reward decoder. Sucrose reward excite neurons had larger normalized beta values overall compared to sucrose reward inhibit neurons (Figure 4H, Mann-Whitney U test: p=0.0002). Taken together, these findings show that sucrose reward excite neurons are bigger drivers of rewarded/unrewarded trial decodability, possibly conveying information about the presence and absence of reward more strongly to downstream regions compared to the sucrose reward inhibited population.

## Discussion

In this study, we investigated the encoding of social and sucrose (food) rewards in the basolateral amygdala. By taking advantage of a two-choice assay in which mice can freely choose between social and sucrose rewards, we found that non-overlapping populations of BLA neurons represent social and food rewards. Interestingly, in the BLA, there is a more equal (males) or slightly stronger (females) representation of sucrose reward in comparison to social reward. This contrasts with what we have observed in the past in the mPFC, where more neurons respond to social reward compared to sucrose reward in both sexes. Additionally, by restricting access to water, we found that BLA neurons are sensitive to internal state changes, with more BLA neurons recruited in response to sucrose reward relative to mice with *ad libitum* access to water. Finally, we found that BLA neurons are sensitive to omissions of expected reward, with BLA neurons responding differentially to rewarded and unrewarded trials upon reward port entry. These findings provide insight on the neural mechanisms underlying social and nonsocial reward choice and consumption within the BLA, as well as highlight the utility of within paradigm comparison across brain regions.

### Social and food reward representations in the basolateral amygdala

The role of the BLA in encoding valence information is well established^33–37^. BLA neurons are known to respond to both positive and negative valence stimuli robustly and have been shown to play an important role in driving approach and avoidance behaviors^44,67,121–124^. While the BLA has been most extensively studied in the context of food rewards^47,72,125,126^ and negative valence stimuli (e.g., foot shocks)^127–134^, more recent work has suggested a key role for the BLA in evaluating social stimuli^67,119,120,135–140^. Because of the highly salient nature of social stimuli and the key role social interactions play in the survival of most species, it has been hypothesized that distinct subpopulations of neurons encode social information^23,74,135^.

In contrast to this idea, one of the first studies to compare social and food reward representations in the BLA involved comparing BLA responses to juice rewards and faces of familiar conspecifics in rhesus macaques^78^. The study found that neurons that were sensitive to juice reward magnitudes were also sensitive to social hierarchy information, suggesting that BLA neurons were multiplexing social and non-social information. However, it should be noted that the social stimuli, in this case, the faces of conspecific monkeys, were used as a conditioned stimulus that was predictive of the subsequent juice rewards, possibly pointing to a role for BLA in learning cue-reward associations, making direct comparisons of reward responses to social and food stimuli difficult in this case.

Conversely, a recent study showed that when rats were serially presented with different conspecifics and foods in a free-range assay, largely different groups of neurons were responsive to each stimulus^79^. However, because animals engage in very different behaviors during social interactions and food consumption during free-range interactions, differences in activity observed during this period could simply be attributed to motor differences. Consistent with this challenge, they found larger differences in BLA neuron activity during contact with the stimuli than during other periods. Therefore, it remains to be determined how the appetitive aspects of social and non-social reward seeking are represented in the BLA. Additionally, this study studied only male rats, and other studies have reported sex differences in social and non-social reward representations in the brain and reward-seeking behavior^25,141^, including the BLA^142–145^, necessitating experiments across both sexes.

The two-choice assay allows us to dissociate the appetitive (choice and approach) and consummatory components of social and food rewards and allows the animals to flexibly and freely choose between these different reward types at the same time. Using the two-choice assay, we found that largely non-overlapping populations of BLA neurons in both female and male mice respond to social and food rewards. This is true both during the choice port entry (Supp. Figure 4D) and reward zone entry (Figure 3D). BLA neurons were specialized in their response to choice and rewards (Supp. Figure 2-3). Additionally, BLA neurons differentially represented social and sucrose rewards at the population level (Figure 3E-G).

Interestingly, a study found that inactivating the BLA decreased female rats’ natural preference for social odors compared to nonsocial ones, with the effect being most pronounced for female urine^146^. However, this manipulation did not disrupt their capacity to tell apart two odors from the same category, such as urine from two different male rats. This finding raises the possibility that the rewards from the same reward category might be similarly represented in the BLA. It would therefore be interesting to determine in the future if more overlapping populations of BLA respond to two different social rewards if animals could flexibly choose between them. For example, mice could choose between two opposite-sex conspecifics or two conspecifics of the same sex (male vs male, or female vs female). In the case of nonsocial rewards, a recent study found that dissociable populations of BLA neurons can represent two different food rewards^72^. Thus raising the intriguing possibility that the BLA processes categorical social information differently compared to food rewards.

### Comparing reward representations in the BLA to other reward nodes in the brain

The BLA is a central node in the reward pathway and is thought to play a crucial role in evaluating and encoding the valence of stimuli^33,34,40,50^, driving approach^41,96,147,148^ and avoidance^6,103,138,149^ behaviors. It projects robustly to the nucleus accumbens^89,96,97,121,150–152^ and receives direct dopaminergic input from the VTA^147,151,153–156^. In addition, it is strongly reciprocally connected to the mPFC^80,81,100,157–159^, another key region that is heavily implicated in both social behaviors^88,135,160–163^ and reward processing^25,83,135,164^.

Interestingly, we found the BLA, like two of its major cortical inputs - the mPFC^25,176^ and OFC^29,165^, has largely non-overlapping social and food reward representations. Similarly, in the NAc, a structure that receives inputs from the BLA, PFC, and the OFC^166–169^, food and social stimuli recruit largely distinct populations of neurons^170^, suggesting that social and food reward-related information remains segregated in most nodes of the reward circuitry. An exception may be the VTA^31^, a region that encodes reward value and reward prediction error, which responds more similarly to social and food stimuli, suggesting that VTA dopamine neurons might purely care about reward value regardless of reward identity.

Although both the mPFC and BLA have largely non-overlapping populations that represent social and food rewards, we found several interesting differences in how these two nodes represent rewards. For instance, the mPFC is more strongly modulated by social reward than by food reward. In contrast, similar proportions of neurons are recruited in the BLA by social and food rewards in males, and in females the bias flips towards sucrose reward (Figure 2E). In both female and male mice, more mPFC neurons are recruited by the social reward than in the BLA. Additionally, unlike the mPFC, choice decisions cannot be decoded above chance from BLA activity (Supp. Figure 4G), suggesting that BLA is more involved in evaluating the reward as opposed to guiding choice decisions. In contrast to the mPFC, significant proportions of BLA neurons are inhibited in response to the social reward. However, it is unclear if these differences point to local microcircuit architectural differences between these two reciprocally connected regions implicated in reward processing.

Previous work has demonstrated sex differences in strategies for learning to attain^171^, motivation to seek^172^, and sensitivity to the outcome of reward^173^. In the mPFC, we found that while female and male mice have similar motivations toward sucrose and social rewards in the two-choice assay, females had more distinct reward representations than males, with stronger responses to social reward compared to sucrose reward, a pattern not seen in males^25^. In the BLA, however, we did not see a difference in the proportion of specialized reward neurons between females and males (Supp. Figure 3). Additionally, we decoded reward type from BLA neural data with similar accuracy between females and males (Figure 3G). These findings suggest that under normal conditions there are only modest sex differences in the representations of food and social rewards. However, during water restriction sex differences in social and sucrose reward became more apparent, with relatively more sucrose reward and less social reward representation in females compared to males (Supp. Figure 6). Water deprivation leads to sex-dependent effects on saline intake and plasma corticosterone levels, with adult females demonstrating higher saline intake and cortisone levels compared to the males^174,175^. Thus, it may be that in the BLA, in contrast to the mPFC, sex-specific representation of various reward types may depend on sex-dependent changes in internal state.

Thus, by comparing neural responses across two different reward nodes in the brain within the same assay, we have been able to gain insight into how the brain represents food and social rewards in both females and males.

### Functional differences in BLA responses to food reward

Evidence suggests that BLA is composed of neurons that are molecularly, anatomically, and functionally heterogeneous^64,125,176–197^. In order to understand how one region can be involved in valence encoding across sensory modalities and contexts, researchers have begun to combine techniques to provide clarity on the functional organization of the BLA. For example, in using a genetic screen one study found two genetically and spatially segregated populations of BLA neurons that encoded either negative valence, Rspo2+ neurons in the anterior BLA, or positive valence, Ppp1rb1+ neurons in the posterior BLA^64^. These two populations were then found to drive defensive or appetitive behaviors through their connections to other sets of genetically defined populations in the central amygdala^33,148^. Many studies have defined specific roles for BLA projections to various cortical and mesolimbic regions. BLA neurons projecting to the nucleus accumbens respond to positive valence stimuli^33,69,148^, and optogenetic activation of BLA-NAc neurons is appetitive^89,95,96,121^, promoting self-stimulation. In contrast, BLA neurons projecting to the ventral hippocampus respond to both negative and positive cues evenly^69^, though optogenetic activation of BLA-vHPC neurons, like BLA-mPFC neurons^103^, increases and inhibition decreases anxiety-like behaviors in mice^198^. Additionally, while the BLA is mostly glutamatergic, recent work^70,199–201^ has begun investigating the role of GABAergic neurons in driving behavior and reward. This continued anatomical^176,184,202,203^ and functional characterization^54,100,204–208^ of the BLA yields a rich picture of how the BLA is involved in valence encoding across such varied behaviors, sensory modalities and contexts.

The BLA’s anatomical and molecular heterogeneity likely underlies the diversity of reward responses evidenced in this study. During the two-choice assay, the BLA had more neurons that are excited by sucrose reward relative to those that were inhibited (Figure 2E, 3B). Rather than similarly respond to the absence of expected sucrose reward, sucrose reward excite and inhibit neurons have differing sensitivities to sucrose omission. We found that 63% of sucrose reward excite neurons dampened their activity, while only 1% of sucrose reward inhibit neurons modulated their response at all during sucrose omission (Figure 4). Additionally, while responses at the population-level did significantly change based on expected reward presence for all sucrose reward responsive neurons, we find that sucrose reward excite neurons may encode more information regarding omissions compared to sucrose inhibit neurons at population level (Figure 4G-H). Whether these neurons route this information to different downstream regions remains unresolved. In our experiments we employed a non-specific GCamP6f virus to image in the BLA, thus it is possible that the patterns we see in reward omission responsiveness, and in social/food responsiveness more broadly, are due to specific cell-type- or projection-specific populations of BLA neurons. These findings provide insight into how food reward, and omission, is processed by the BLA, enabling future studies to explore how different circuits and/or cell-types may give rise to this pattern.

### What about social reward - different valence stimuli and social reward omissions

Through this study, we found that the BLA distinctly represents sucrose and social reward in both baseline and water deprived conditions. Our results pointing to a diversity of reward omission responses in sucrose reward neurons prompts whether social reward responsive neurons may also respond differentially to social reward omission. This exploration may require modification of the existing two-choice assay, or a new paradigm optimized for these questions. Given the complexity of social stimuli^209–213^, investigating this question requires the operationalization of a “social reward omission”. Does social reward omission require the elimination of auditory, olfactory and/or tactile engagement? Investigating which sensory modalities, or in what combination, has the largest effect on BLA activity would be important and necessary to interpret findings. Nevertheless, if social reward omission yields similar patterns in social reward neurons as evidenced in sucrose reward neurons, it could point to a shared neural mechanism for the registering of expected reward outcomes across reward types.

We focus this study on how the BLA differentially responds to positively valenced rewards, sucrose consumption and novel same-sex interaction. However, given the role of the BLA in assessing both positively and negatively valence stimuli^33–41^, it would be interesting to explore how the BLA may encode social “reward” with neutral or negatively valenced social targets. Or how representations may shift when the relative value of the social and food reward shifts. Recent work has shown that BLA neurons quickly re-scale their value representations of food rewards when a new, higher value food is presented^72^. It has yet to be explored if BLA neurons would also shift their relative representations when rewards of different modalities (i.e. social vs nonsocial) shift from their original value. Such experiments could provide insight into how the brain processes and guides decisions to seek or avoid various stimuli in an animal’s environment.

### Conclusions and future directions

Our findings reveal that distinct populations of BLA neurons respond robustly to social and food (sucrose) rewards. In contrast to our previous work in the mPFC, BLA neurons did not exhibit a bias towards social reward, and instead demonstrated either similar social/sucrose representations, the case in males, or a bias towards sucrose reward, the case in females. Water deprivation increased sucrose reward representations and strengthened sex differences in neural responses. Lastly, sucrose omission revealed functionally distinct groups of sucrose reward neurons, where sucrose reward-excited neurons were more sensitive to expected sucrose reward omissions compared to neurons that were originally inhibited by reward. This study adds to a growing body of work in the BLA that aims to elucidate whether the BLA uses a shared, common currency population or distinct populations to process various stimuli. While this work supports the latter, it may be important to consider agency in how the brain may employ various neural populations to encode valence and salience of stimuli. Whether an animal has the true choice to engage with a social or nonsocial stimulus could very well affect how these stimuli are processed. Systematically comparing how different brain regions, like the BLA and mPFC, encode reward-related information within the framework of the same task allows us to identify differences and similarities in how the brain processes social and non-social information and ultimately helps shed light on how these nodes function in the context of reward-related decision making.

## Supplementary Figures

**Supplementary Figure 1:**
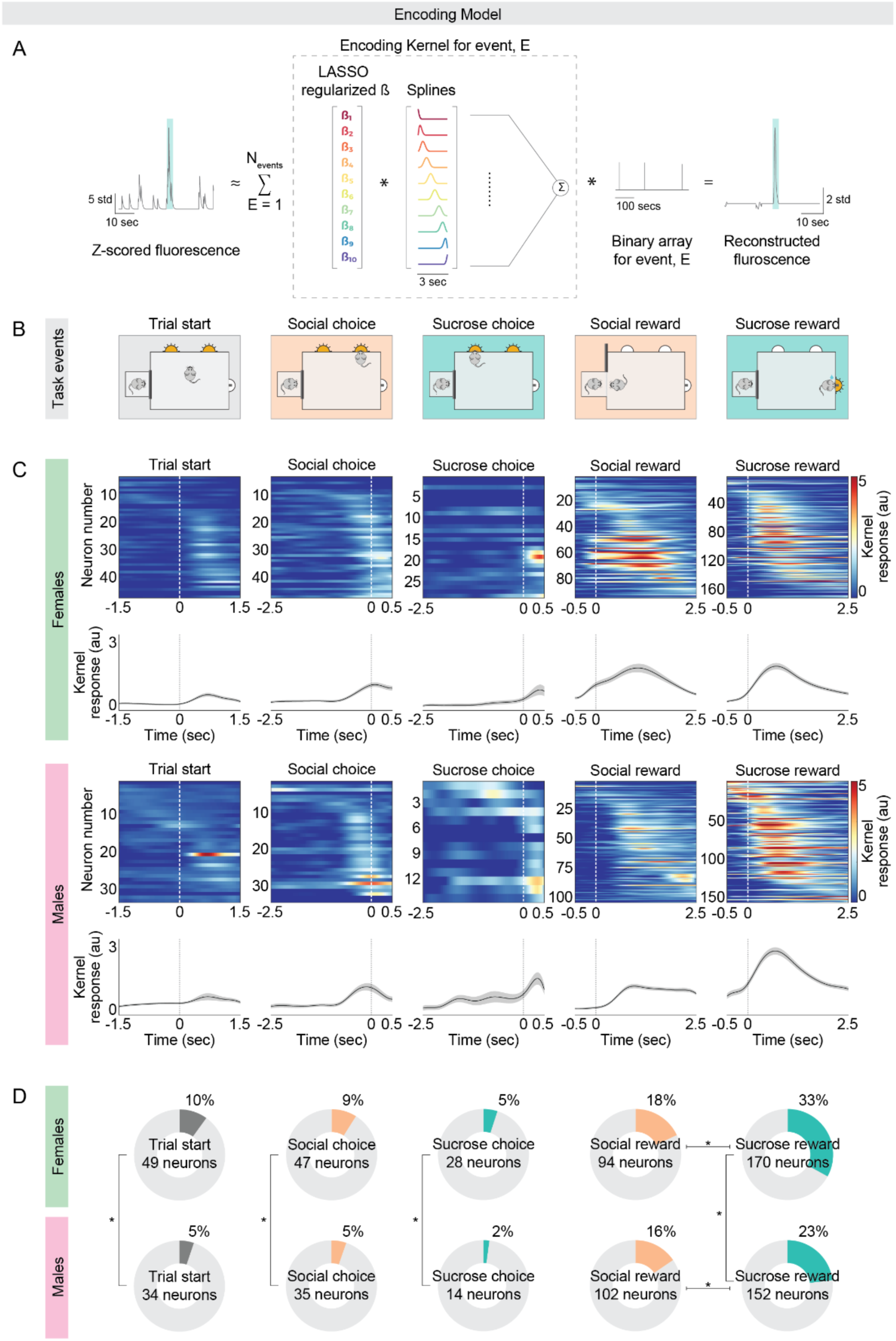
An encoding model identifies similar patterns of task-modulated neurons in the BLA. A. Schematic of the encoding model adapted from Isaac et al 2024. A response kernel was created for each neuron for each operant task event. For each neuron, a fluorescence trace was estimated as the sum of response kernels for all events, convolved with a binary array for each event. A reconstructed fluorescence for one example neuron during one trial of an event is shown on the right. B. Two-choice operant assay task events. C. Heatmaps are of average kernel responses (au) from BLA neurons significantly modulated by each task event. Neurons are sorted by the time of maximum response across each event. A dashed line at zero shows task event onset. Below the heatmaps are plots of the average kernel response traces (au) of the neurons that are significantly modulated by each task event from the corresponding heatmap. Shaded error regions indicate ± SEM. D. In donut charts are the proportions of total BLA neurons that are modulated by each task event in females and males as determined by the encoding model (female: n=513 neurons, 7 mice; male: n=658 neurons, 7 mice). Females show higher representations of trial start, social choice, sucrose choice and sucrose reward, but not social reward, compared to males (proportion z-test, female vs male: trial start: p=0.0037, social choice: p=0.0105, sucrose choice: p=0.0024, social reward: p=0.2005, sucrose reward: p=0.0001). Both females and males had proportionately more sucrose reward representation compared to social reward (proportion z-test, social reward vs sucrose reward, female: p<0.00001; male: p=0.0005). The Bonferroni corrected threshold for proportion comparisons for social and sucrose reward it’s p=0.025 (female vs male, social vs sucrose), for trial start, social and sucrose choice is p=0.05 (female vs male).

**Supplementary Figure 2:**
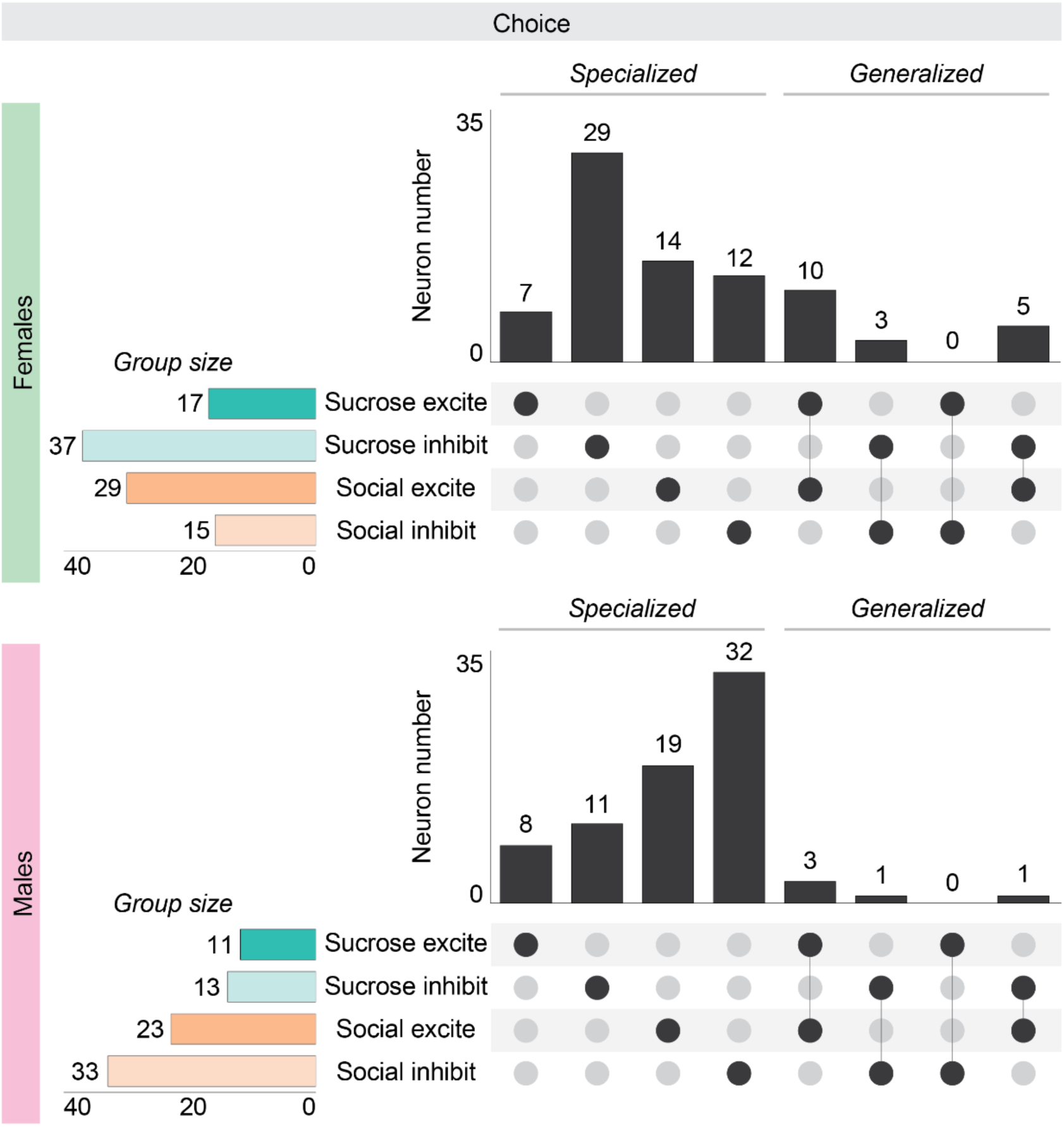
The BLA neurons are selective in their choice responses to sucrose and social reward. UpSet plots showing the membership of individual choice responsive neurons to different response groups for female and male mice. Response group size is counted as the sum of all filled in circles horizontally. Vertical bars represent the number of neurons defined by the filled in circles directly below. Connected lines mean neurons are defined by membership to both response groups. Overlap between choice response types for significantly responsive neurons in females and males. Overall, most neurons only responded to either sucrose or social choice in both females and males (proportion z-test: female: specialized: 77.5%, generalized: 22.5%, n=80 neurons, p<0.00001; male: specialized: 93.3%, generalized: 6.67%, n=75 neurons, p<0.00001). Males had proportionately more specialized choice neurons compared to females (proportion z-test: female specialized 62/80 vs male specialized 70/75, p=0.0056). The Bonferroni corrected threshold for proportion comparisons for specialized neurons is p=0.025 (specialized vs generalized, female vs male).

**Supplementary Figure 3:**
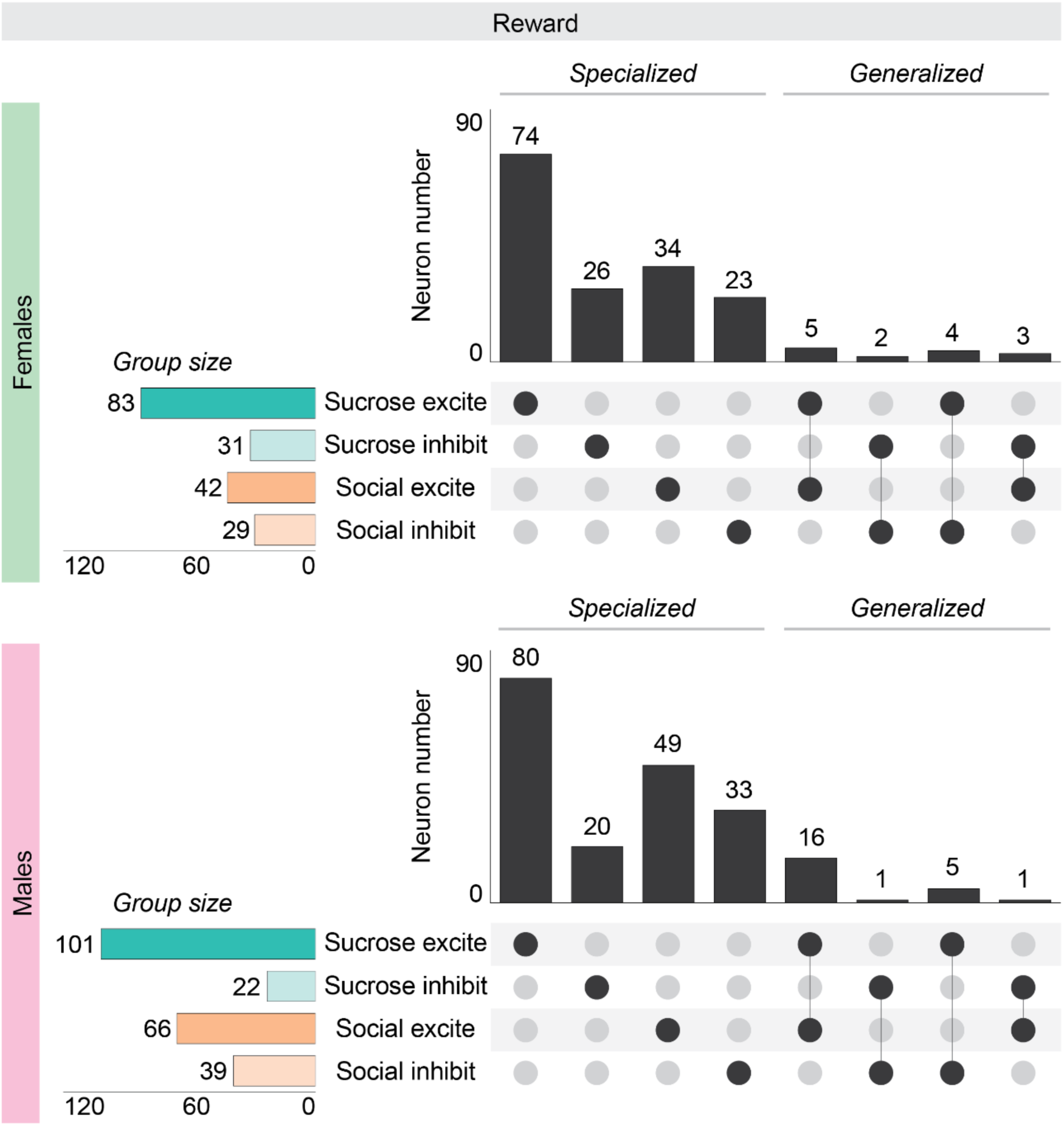
BLA neurons often respond selectively to either sucrose or social reward. UpSet plots showing the membership of individual reward responsive neurons to different response groups for female and male mice. Response group size is counted as the sum of all filled in circles horizontally. Vertical bars represent the number of neurons defined by the filled in circles directly below. Connected lines mean neurons are defined by membership to both response groups. Overlap between reward response types for significantly responsive neurons in females and males. Like during choice, most neurons only responded to one type of reward in both females and males (proportion z-test: female: specialized: 88.8%, generalized: 11.2%, n=171 neurons, p<0.00001; male: specialized: 93.3%, generalized: 6.67%, n=205 neurons, p<0.00001). Females and males had similar proportions of specialized reward neurons (proportion z-test: female specialized 157/171 vs male specialized 182/205, p=0.0056). The Bonferroni corrected threshold for proportion comparisons for specialized neurons is p=0.025 (specialized vs generalized, female vs male)

**Supplementary Figure 4:**
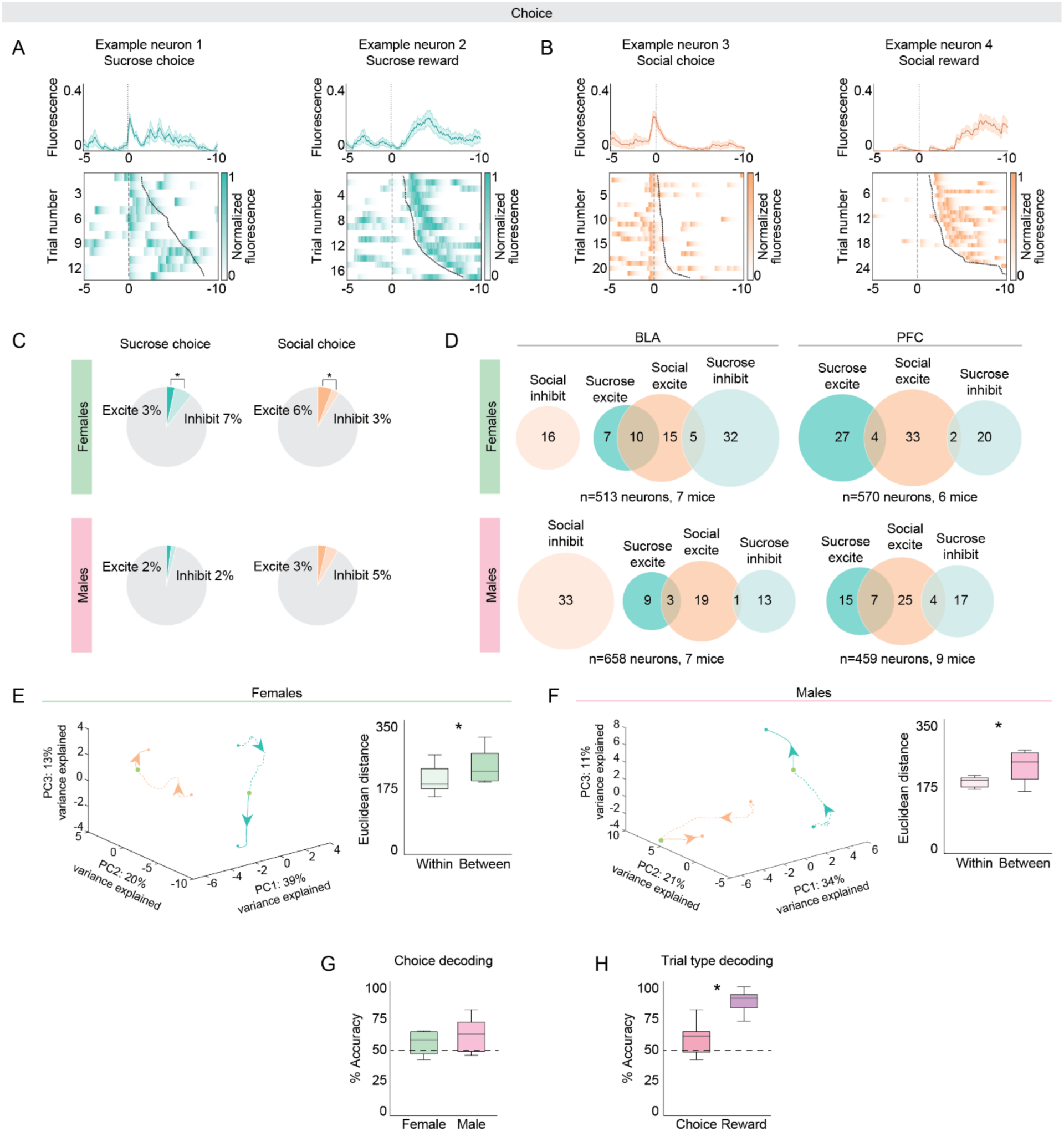
Sucrose and social choice representations are mostly non-overlapping but less decodable than reward representations. A. Sucrose choice responses are separable from sucrose reward responses. The top row shows a normalized fluorescence trace of one example neuron that is significantly tuned to sucrose choice (left) and a different neuron significantly tuned to sucrose reward (right). A dashed line at 0 shows the onset of sucrose choice. The bottom row displays heatmaps of the activity over trials of the corresponding neuron. In both heatmaps, the dashed line at 0 is aligned to the onset of sucrose choice. The solid black line shows the onset of sucrose reward. Trials are sorted in ascending order of reward latency. B. Similarly, social choice responses are separable from social reward responses. Example neuron 3 (left) is significantly tuned to social choice and example neuron 4 (right) to social reward. C. Proportions of total recorded BLA neurons that are significantly responsive to sucrose choice and social choice in females and males (female: n=513 neurons, 7 mice; male=658 neurons, 7 mice). In females, there are proportionately more neurons inhibited rather than excited during sucrose choice (proportion z-test: sucrose excite: 17/513, sucrose inhibit: 37/513, p=0.0010). Conversely, females had more neurons excited as opposed to inhibited by social choice (proportion z-test: social excite: 30/513, social inhibit: 16/513, p=0.0349). Males had similar proportions of choice excite and inhibit neurons for both sucrose and social trials (proportion z test: sucrose excite: 12/658, sucrose inhibit: 14/658, p=0.6892; social excite: 23/658, social inhibit: 33/658, p=0.1707). D. Venn diagrams displaying the overlap of neurons that are either excited or inhibited by sucrose and social choice in the BLA and mPFC. Circle sizes are proportional to the number of neurons in each group. Proportions of mPFC neurons for comparison in each respective group from Isaac et al (2024). Not shown: BLA overlap between social inhibit and sucrose inhibit for females is 3 neurons, for males is 1 neuron. E. Left: Principal component analysis of trial-averaged population neural activity for both sucrose (teal) and social (orange) choice trials for females (n=513 neurons). Arrowhead indicates direction of time. Filled green circle indicates choice onset. Right: The Euclidean distance separating the PC-projected population vectors in the choice window is significantly greater between social and sucrose trials than within each trial type (unpaired two sample t-test, within vs between, p<0.00001). Shaded error regions indicate +/− SEM. F. Same as E, left: trial-averaged population traces for sucrose and social choice trials plotted on the first 3 principal components in state space for males (n=658 neurons). Right: Euclidean distance between social and sucrose trial population vectors was greater than within trial type (unpaired two sample t-test, within vs between, p=0.0027). G. Decoders trained on neural data were unable to distinguish between social and sucrose choice in both male and female mice (unpaired two sample t-test, female vs male, p=0.3038; paired two sample t-test, female vs shuffled, p=0.4542, male vs shuffled, p=0.0824). Decoding accuracy was calculated for each animal from all recorded neurons on the test set (70-30 train-test data), with a trial-matched number of sucrose and social trials. H. When trained on both female and male neural data from the choice time window, a decoder was unable to distinguish between sucrose and social trials (paired two sample t-test, choice vs shuffled, p=0.0538). A decoder trained on both female and male neural data from the reward time window, however, was able to correctly identify sucrose and social trials above chance (paired two sample t-test, choice vs shuffled, p<0.00001, reward vs choice, p<0.00001). Box plot center line indicates the median, box edges indicate the 25th and 75th percentile and whiskers extend to +/− 2.7x the standard deviation. N=7 female mice, 7 male mice.

**Supplementary Figure 5:**
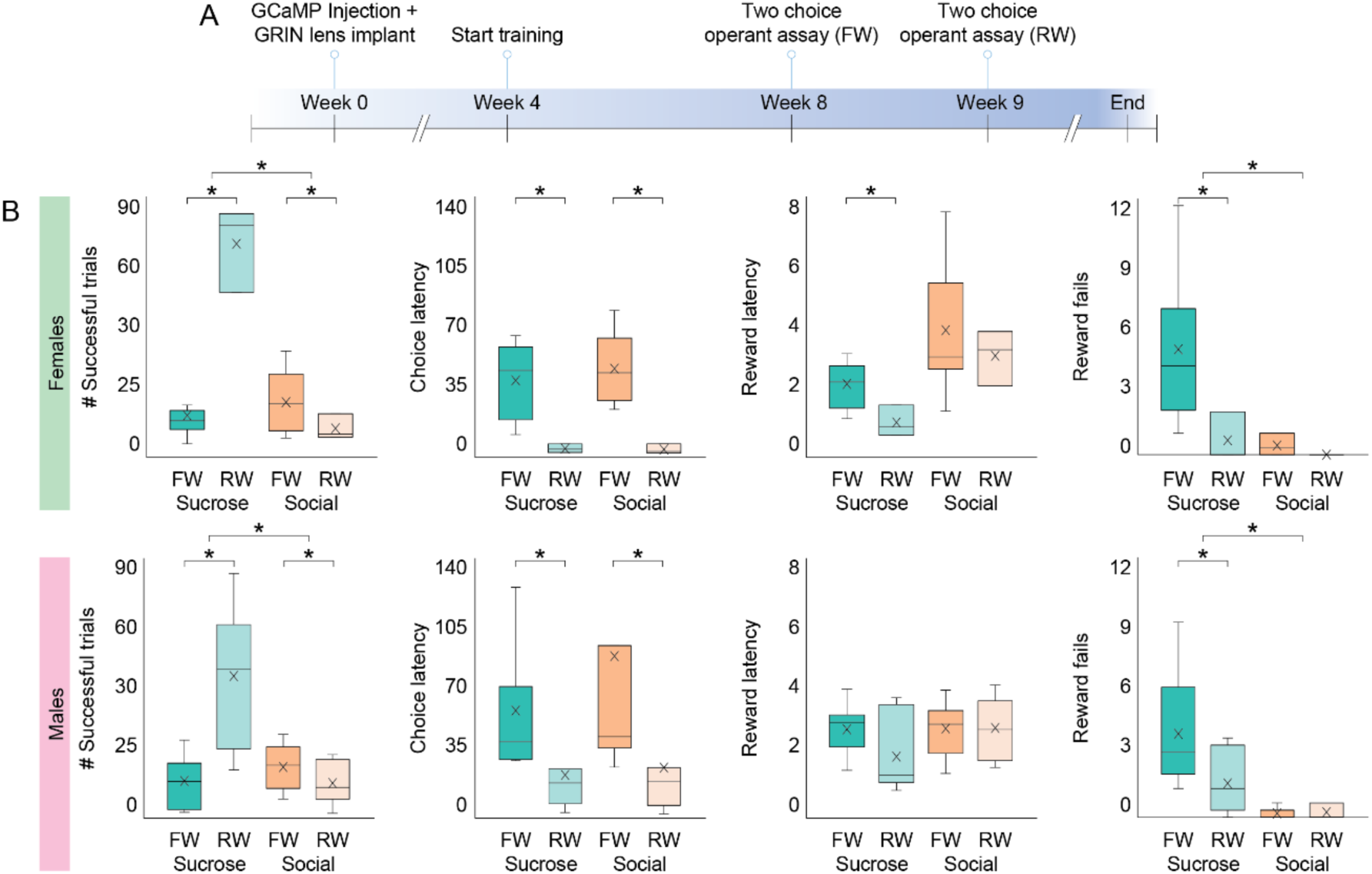
Mice increase their sucrose reward seeking during restricted water access. A. Timeline of the experimental schedule. Mice are trained and then go through two-choice testing with full (*ad libitum*) access to water (FW) then are water restricted and go through two-choice testing again (RW). B. During restricted water access both females and males increase their successful sucrose trials and decrease their successful social trials (two-factor anova, factors: water condition (FW/RW), trial type (sucrose/social): female: interaction p<0.00001, water condition p<0.00001, trial type p<0.00001; male: interaction p<0.00001, water condition p<0.00001, trial type p<0.00001; post-hoc unpaired t-test comparing FW/RW within trial type: female: sucrose: p<0.00001, social: p=0.0013; male: sucrose: p<0.00001, social: p=0.0466). Both females and males displayed generally decreased choice latencies during RW compared to FW (two-factor anova, factors: water condition (FW/RW), trial type (sucrose/social): female: interaction p=0.8079, water condition p<0.00001, trial type p=0.3938; male: interaction p=0.257, water condition p=0.0007, trial type p=0.1495; post-hoc unpaired t-test comparing FW/RW within trial type: female: sucrose: p<0.00001, social: p=0.0003; male: sucrose: p=0.0035, social: p=0.0136). Females had decreased sucrose reward latencies during RW, while males showed no difference across water conditions or trial types (two-factor anova, factors: water condition (FW/RW), trial type (sucrose/social): female: interaction p=0.6169, water condition p=0.0038, trial type p<0.00001; male: interaction p=0.1395, water condition p=0.1621, trial type p=0.1395; post-hoc unpaired t-test comparing FW/RW within trial type: female: sucrose: p<0.00001, social: p=0.2143; male: sucrose: p=0.0333, social: p=0.9569). Both females and males displayed decreased sucrose reward fails during RW and overall more sucrose reward fails compared to social reward regardless of water condition (two-factor anova, factors: water condition (FW/RW), trial type (sucrose/social): female: interaction p=0.004, water condition p<0.00001, trial type p<0.00001; male: interaction p=0.0345, water condition p=0.062, trial type p<0.00001; post-hoc unpaired t-test comparing FW/RW within trial type: female: sucrose: p<0.00001, social: p=0.0003; male: sucrose: p=0.0333, social: p=0.7706).

**Supplementary Figure 6:**
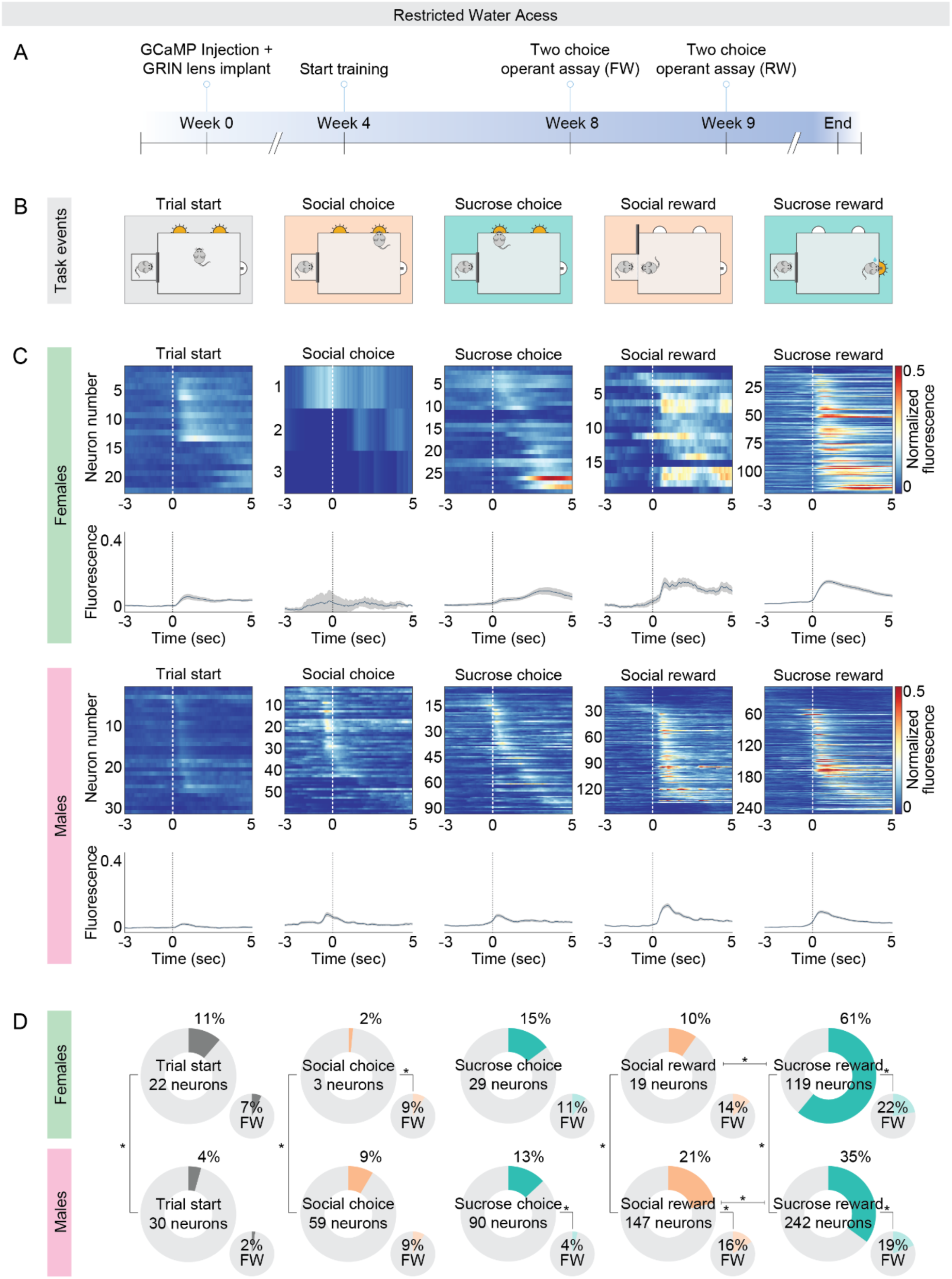
Increased sucrose reward representations in response to water restriction in both male and female mice. A. Experimental timeline. Mice are trained, go through two-choice testing with full (ad libitum) access to water (FW), then are water deprived and placed under restricted access to water (RW). B. Schematic of two-choice operant assay task events to which BLA activity was aligned, from left to right: trial start, social choice, sucrose choice, social reward, sucrose reward. C. Top row: Neuronal data for significantly modulated neurons in females and males. For both females and males, heatmaps are of average fluorescence across trials of neurons that are significantly modulated by task events. Neurons are sorted by the time of maximum fluorescence across each task event. Neurons that are classified as modulated by more than one task event are included in all relevant heatmaps. Bottom row: Average fluorescence traces of the neurons that are significantly modulated by each task event from the corresponding heatmap. Shaded error regions indicate ± SEM. D. In large donut charts are the proportions of total recorded BLA neurons that are modulated by each task event in females and males during restricted water access (female: n=195 neurons, 3 mice; male: n=691 neurons, 6 mice). In the small pie charts are the proportions of BLA neurons that were modulated by each task event during full (*ad libitum*) access to water (FW). Females and males represent task parameters differently during restricted access to water. Females show a greater proportion of trial start and sucrose reward responses, a lower proportion of social choice and social reward responses, and similar proportion of sucrose choice responses compared to males (proportion z-test, female vs male: trial start: p=0.00028, social choice: p=0.00072, sucrose choice: p=0.50286, social reward: p=0.00028, sucrose reward: p<0.00001). However, both females and males more highly represent sucrose reward compared to social reward during restricted water access (proportion z-test, female: p<0.00001, male: p<0.00001). Compared to FW, females during RW showed decreased social choice response and an increased proportion of sucrose reward response (proportion z-test, RW vs FW: trial start: p=0.06432, social choice: p=0.0005, sucrose choice: p=0.1074, social reward: p=0.1443, sucrose reward: p<0.00001). Compared to FW, males during RW demonstrated an increase in the proportion of sucrose choice response and sucrose reward response neurons, and a decrease in social reward response neurons (proportion z-test, RW vs FW: trial start: p=0.0536, social choice: p=0.98404, sucrose choice: p<0.00001, social reward: p=0.01242, sucrose reward: p<0.00001). Schematic in A adapted from Isaac et al. (2024). The Bonferroni corrected threshold for proportion comparisons for trial start, social and sucrose choice is p=0.025 (female vs male, BLA vs mPFC), for social and sucrose reward it’s p=0.017 (female vs male, BLA vs mPFC, social vs sucrose).

**Supplementary Figure 7:**
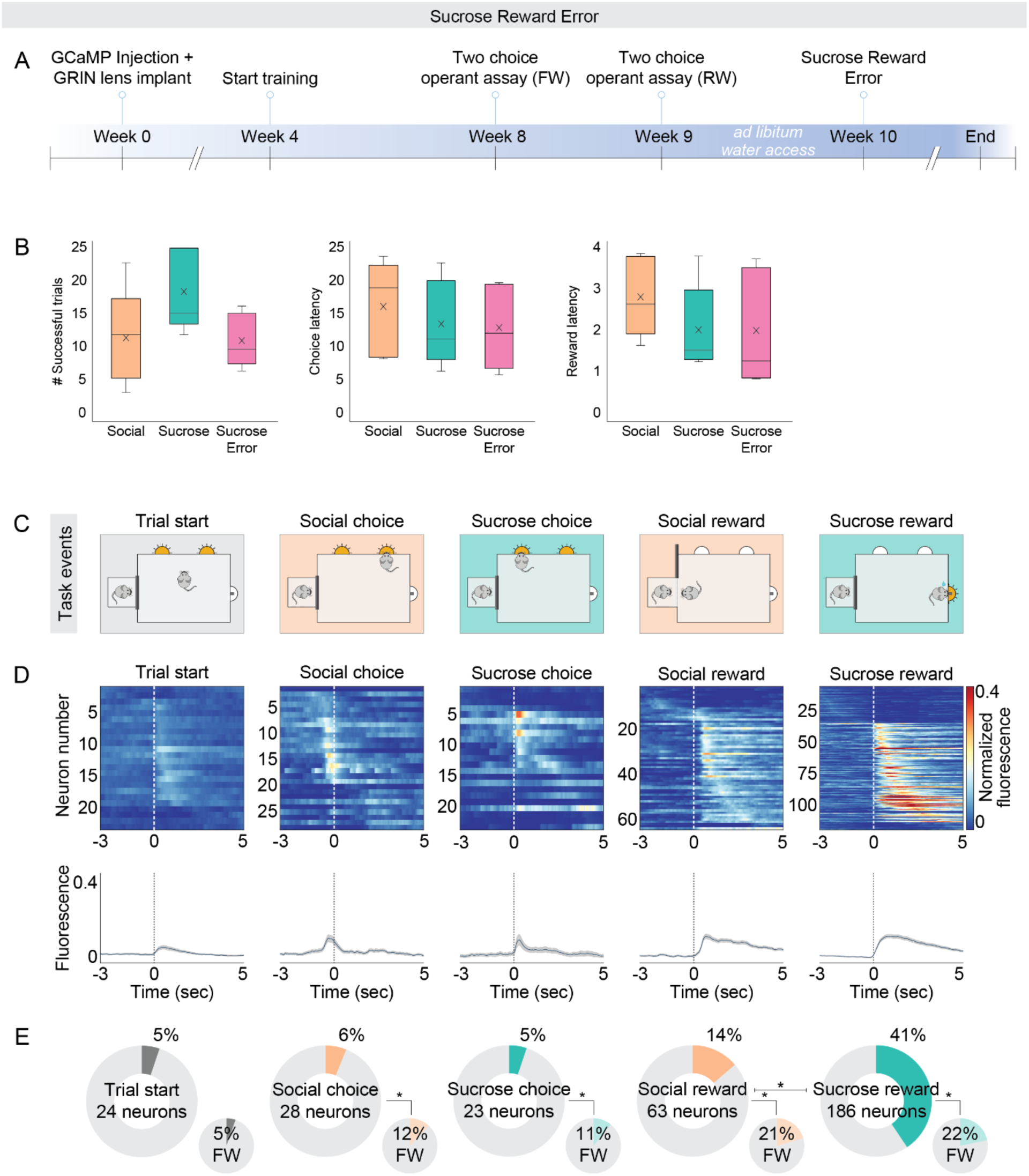
Behavioral tracking and neural responses during sucrose reward omission. A. Timeline of the experimental schedule. Mice are trained and then go through two-choice testing with full (*ad libitum*) access to water (FW), they’re then water deprived and go through two-choice testing with restricted access to water (RW). Mice are then again given *ad libitum* water access before undergoing one session of a modified two-choice assay where 30% of sucrose rewards are randomly withheld. B. Behavioral results from this modified sucrose omission two-choice assay (n=5 mice, 3 female, 2 male). Mice display typical two-choice behavior even when the sucrose reward becomes unpredictable. C. Two-choice operant assay task events. D. Neuronal data for significantly modulated neurons. Heatmaps display average normalized fluorescence of neurons across individuals that are significantly modulated by task events. Neurons are sorted by the time of maximum fluorescence across each task event. Neurons that are classified as modulated by more than one task event are included in all relevant heatmaps. Below heatmaps are plots of the average normalized fluorescence traces of the neurons that are significantly modulated by each task event from the corresponding heatmap. Shaded error regions indicate ± SEM. E. In large donut charts are the proportions of total recorded basolateral amygdala neurons that are modulated by each task event in females and males (n=454 neurons, 5 mice). In the small pie charts are the proportions of BLA neurons, in the same mice, that were modulated by each task event during full (*ad libitum*) access to water when no omissions occurred (FW, n=461 neurons, 5 mice). During the sucrose omission session, mice had decreased representations of social and sucrose choice, and social reward compared to the full water access condition but increased sucrose reward representation (proportion z-test, trial start: p= 0.9045, social choice: p=0.0021, sucrose choice: p<0.00001, social reward: p=0.0060, sucrose reward: p<0.00001). Additionally, there was a larger fraction of sucrose reward responsive neurons compared to social reward during the sucrose omission session (proportion z-test, p<0.00001). The Bonferroni corrected threshold for proportion comparisons for social and sucrose reward is p=0.025 (sucrose omission vs FW, social vs sucrose), for trial start, social and sucrose choice is p=0.05 (sucrose omission vs FW).

## Methods

### Mice

All mice involved in this study were C57BL/6J (Jackson Laboratories) female and male mice aged between 6-10 weeks at the start of the experiment. Mice were maintained on a reverse light dark cycle (12h light off - 12h light on schedule) during all experiments. All experiments were performed during their night cycle. Mouse sex was defined by external anatomy. All experimental procedures were approved by the Emory Institutional Animal Care and Use Committee.

### Stereotaxic Surgery

Mice aged ∼6-10 weeks were anesthetized with 1–2% isoflurane and placed in stereotactic setup (Kopf). A Nanoject microsyringe was used to inject 750 nL of AAV5-Syn-GCaMP6f-WPRE-SV40 (100835-AAV5, Addgene, injected titer of 1.3 × 10^13^ parts/ml) unilaterally into the BLA (AP −1.6 mm, ML +/− 3.4 mm, DV −4.8 mm). Then, mice were implanted with a 0.6 mm diameter, 7.3 mm length GRIN lens (Product ID:1050-004597, Inscopix) over the BLA (AP −1.6, ML +/− 3.4, DV - 4.9). A metal cap was cemented over the lens to protect it. Mice were group housed following lens implant. 3–4 weeks following viral injection and lens implant, a baseplate (Product ID: 1050-004638, Inscopix) attached to the miniature microscope (nVue, Inscopix) was positioned ∼0.45 mm above the lens. Blood vessels and neurons were used as landmarks to refine the focus of the lens. The baseplate was then cemented in place using dental cement (Metabond) and a baseplate cover (custom design) was secured over the baseplate to protect the lens.

### Automated Operant Assay

The two-choice operant assay was conducted as previously described; in depth, complete details on the building and execution of the two-choice operant assay can be found in Isaac & Balasubramanian (2025) and Isaac et. al (2024).

Briefly, mice were placed in an acrylic chamber (12×12×12 inches) outfitted with two-choice ports, a reward port and a social reward access zone. During the two-choice assay, the beginning of a trial illuminates both choice ports via LED. Animals are completely free to choose between either choice port. When a nosepoke is registered via an infrared beam break at the sucrose choice port, it triggers LED illumination of the sucrose reward port. Once an infrared beam break is registered at the sucrose reward port, a solenoid valve connected to a liquid reservoir releases 10 µL of 10% sucrose/water solution. Sucrose reward consumption must happen within 8 sec of sucrose choice to count as a successful trial, or the trial will end. If at the beginning of a trial, a nosepoke is registered at the social choice port, then a gate will open allowing the experimental mouse 20 sec of social interaction with a novel same-sex conspecific through a set of bars, preventing entry into the social targets chamber. At the end of 20 sec, the social trial is over and the gate closes. Each trial is separated by a 30 sec intertrial interval, after which a new trial begins. Each behavioral session is 1 hour. All ports and the social reward gate were operated using Sanworks Bpod.

Each mouse is trained in four training stages as explained in Isaac & Balasubramanian (2025). After training, all mice were tested on the two-choice operant assay with full *ad libitum* water access for 5 days, 1 session each day (n=10 female mice, 7 male mice).

#### Restricted Water Access

Following testing on the two-choice operant assay with full water access, a subset of animals underwent testing in two-choice operant assay with restricted water access. Mice were first placed on a restricted water schedule for 2-3 days prior to testing, receiving 1 mL of water each day. Mice were then tested in the two-choice operant assay for 5 days, 1 session each day (n=3 female mice, 6 male mice). The weight of each mouse was monitored each day to ensure mice never fell below 85% of their starting weight.

#### Sucrose Omission

After 5 days of the two-choice operant assay with restricted water access, a subset of the mice were placed on full water access for 2-3 days. These mice were then tested on a modified version of the two-choice operant assay during a single session wherein on ∼30% of sucrose trials sucrose reward was omitted (n=3 female mice, 2 male mice).

### Behavioral Data Analysis

Videos for each behavioral session were recorded at 40 Hz using Pylon software. Mouse behavior was quantified during each session by the following metrics: number of successful pokes, choice latency, reward latency and reward fails. Successful pokes were defined as the number of trials when a mouse successfully made a choice and recruited the respective reward. Choice latency is defined as the time between trial start (choice port LEDs turn on) and a nosepoke at a choice port. Reward latency is defined as the time between a nosepoke at a choice port and recruitment of the respective reward. For sucrose trials, reward recruitment is defined at nosepoking at the sucrose reward port after sucrose choice. For social trials, reward recruitment is defined by entry into the social reward zone, a 1×3 inch area in front of the social gate after social choice. Reward fails are trials when a mouse successfully makes a choice but fails to recruit the respective reward by either not nosepoking in the sucrose reward port within 8 sec of sucrose choice or not entering the social reward zone within 20 sec of social choice.

To quantify the described metrics, LED light levels and mouse body position (centroid) were tracked and analyzed using Bonsai software^214^. Using the body position data, we then identified when a mouse entered the sucrose or social reward zone. With tracked LED light levels for each choice port, we identified the time of trial start and choice. Data collected and analyzed with Bonsai was combined with session data generated by Bpod to quantify reward behavior for each mouse, for each behavioral session during training and post-training testing. For the full water and restricted water conditions, 5 behavioral sessions were recorded, the last three sessions were considered per mouse in the calculation of behavioral metrics (Figure 1B, Supp. Figure 5B). For the sucrose omission condition, behavior from a single session was used in the calculation of behavioral metrics (Supp. Figure 7).

### Histological Analysis of Lens Placement

Once experiments were completed, mice were sacrificed and perfused with 0.5x phosphate-buffer saline (PBS) followed by 4% paraformaldehyde (PFA). Brains were subsequently removed and fixed in 4% PFA for 1-2 nights at 4 °F. The brains were then placed in 30% sucrose solution for at least 24 hours, after which they were sliced at 50 µm using a microtome. Slices containing the BLA were mounted in DAPI solution (Product ID: 0100-20, Southern Biotech) and imaged using the Keyence BZ-X800. Using the Allen Mouse Brain Common Coordinate Framework^215^, fluorescent images were then used to determine GRIN lens placement. Animals with no to low expression of GCaMP6f or mistargeted lens placements were excluded from calcium imaging analysis (n=3 female mice).

### One Photon Calcium Imaging Experiments and Analysis

On all non-imaging days, from training to testing, mice were habituated to wired miniscope recording using a tethered dummy scope (Product ID: 1050-003762, Inscopix). During the behavioral imaging session, the LED power was set to 0.6 and the analog gain on the image sensor was set to 1.6–1.8. One photon calcium imaging recording was acquired at 20 Hz using Inscopix nVue software. Once recorded, calcium imaging videos were pre-processed using Inscopix software, first spatially downsampled by a factor of 4, then linearly motion corrected or non-linearly motion corrected using NormCorr^216^. Using a CNMFe algorithm^217^, individual neurons were then identified along with their normalized fluorescence traces. Fluorescence traces were then aligned to behavioral data for task modulation analysis. Custom MATLAB code was used for all analyses following CNMFe.

### Data Analysis and Statistics

Analysis for task modulation, development of the encoding model, production of neural trajectories for choice and reward, and construction of neural decoders for choice and reward were conducted as described in detail in Isaac et al (2024).

### Task Modulation Analysis

Using previously described methods^25,218^, neurons were classified as significantly modulated by task events during the two-choice operant assay: trial start, social and sucrose choice, social and sucrose reward. A 3 sec window relative to task event onset was defined: trial start: 1.5 sec before and after; choice: 2.5 sec before, 0.5 sec after; reward: 0.5 sec before, 2.5 sec after. Within each window, the change in fluorescence was analyzed. To identify significantly excited responses, we compared the maximum value of the mean fluorescence to the maximum value of a shuffled distribution (>99.5 percentile). To identify significantly inhibited responses, we compared the minimum value of the mean fluorescence to the minimum value of a shuffled distribution (<0.5 percentile). Shuffled distributions were created by applying a randomized *circshift* (MATLAB) to the individual fluorescence traces of each neuron for 1000 iterations and then averaging shuffled activity within each 3 sec task event-specific time window.

Through this analysis, neurons could be part of multiple task-responsive groups, such that a sucrose excite neuron could also be a social inhibit neuron. We visualized the overlap between response groups, social inhibit, sucrose excite, social excite, sucrose inhibit, for choice and reward responsive neurons using Venn diagrams where circle sizes were proportional to the number of neurons in each group (Supp. Figure 4D, Figure 3B). To clearly demonstrate the intersections between all four response groups, we used UpSet^219^ plots for both choice and reward responsive neurons (Supp. Figure 2-3).

To assess whether individual sucrose reward excite neurons were significantly modulated by the absence of reward (Figure 4A), for each neuron we compared the maximum fluorescence values during sucrose reward trials to the maximum fluorescence values during unrewarded sucrose trials, where sucrose reward was withheld, using an unpaired t-test. Similarly, to identify which sucrose reward inhibit neurons were significantly modulated by the absence of reward (Figure 4D), we compared the minimum fluorescence values during sucrose reward trials to the minimum fluorescence values during unrewarded sucrose trials using an unpaired t-test. To assess population level response differences to rewarded and unrewarded trials, we compared the average fluorescence trace during the reward window (−0.5s to 2.5s) between trial types using a paired t-test (Figure 4B-C, E-F).

### Encoding model

As previously described^25,172^, we used a linear encoding model to estimate neural response kernels for each neuron to identify the contribution of each task event (trial start, social choice, sucrose choice, social reward, sucrose reward) to the variability in fluorescence (Supp. Figure 1). The model included z-scored fluorescence activity of each neuron as the dependent variable and the task events as the independent variable. The task events were configured as a set of binary arrays of event time (1 the event occurs, 0 it does not), convolved with an spline basis set (with 10 splines) spanning 3 s. The independent variables of the multiple-linear regression model capture time-delayed relationships between the task event and the corresponding fluorescence.

The encoding model was then used to identify neurons significantly modulated by task events. We compared the goodness of fit of our encoding model to the fluorescence activity when all events were included to the goodness of fit of a reduced model where one of the events is removed. Goodness of fit for the models was measured using the F-statistic, which was iteratively calculated after removing predictors of each event from the model. F statistics were calculated for both experimental data and for shuffled data, generated by shuffling the fluorescence data in 1-s bins (to preserve correlational structure of calcium data) 500 times. F-statistics were then compared between real and shuffled data. Neurons were considered significantly modulated by a task event when the resulting calculated p-value was less than 0.01 and corrected for multiple comparisons.

### Neural trajectories for choice and reward

Population activity trajectories for sucrose and social trials were constructed using all recorded neurons across trials and reducing them to a three-dimensional neural subspace using principal component analysis (PCA)^220^. To visualize female and male choice (Supp. Figure 4 E-F) and reward (Figure 3 C-D) trajectories, trial-averaged z-scored fluorescence during choice or reward time windows, respectively, were calculated and concatenated across animals to form a three-dimensional array, [number of neurons x trial type x time window]. A PCA was fit onto the array after collapsing the last two dimensions. The resulting dimensionality-reduced array was then projected independently onto social and sucrose trials (score output from *pca*, MATLAB).

We then quantified the difference between the neural space occupied by social and sucrose trials. We repeated the PCA procedure above, this time for each animal, and calculated the pairwise Euclidean distance within and between the first three PC projected vectors of social and sucrose trials.

### Neural decoders for choice and reward

Using logistic regression (*fitclinear*, MATLAB) for neural data during the choice or reward time window, we decoded the identity (sucrose or social) of choice (Supp. Figure 4G) and reward (Figure 3E) trials^25,221^. For each mouse, a three-dimensional array of [number of neurons × number of trials × time window of z-scored fluorescence data] was created using trial-matched numbers of sucrose and social trials. Trial-matching allowed each binary decoder to be trained with balanced labels despite the variability in the number of sucrose and social trials across mice. To maximize the number of samples that could be used to train our decoder, the 3D array was collapsed across the last two dimensions and used as an input to the regression. This input was split into a 70% train and 30% test set. The train set was used to optimize the regression model hyperparameters using Lasso regularization.

To evaluate the performance of the decoder, written as decoding accuracy, we applied the trained model to the test set and evaluated correctly predicted labels (social or sucrose). Decoding accuracy was calculated for 500 iterations of train and test sets created from randomly sampled trials. Decoding accuracies for all 500 iterations were then averaged per mouse, yielding the average decoding accuracy. We then calculated the decoding accuracy of the z-scored fluorescence data shuffled in non-overlapping 1 sec time bins, maintaining the autocorrelation property of the signal while disrupting the activity-aligned neural responses. We then were able to compare decoding accuracies for real vs shuffled data.

A similar process was performed for designing decoders to identify neural responses from rewarded and unrewarded sucrose trials (Figure 4G). In this case, for each mouse, three-dimensional arrays were created using a trial-matched number of rewarded and unrewarded sucrose trials. Decoders were based on neural data from either all neurons or just sucrose reward neurons. To compare the beta values of individual sucrose reward excite and sucrose reward inhibit neurons, we normalized the absolute value magnitudes of the betas to the largest magnitude beta value across all sucrose reward neurons, both inhibit and excite.

### Statistical analyses

All statistical tests were done with custom MATLAB scripts. Unless noted, all statistical tests were two-sided, and Bonferroni corrected for multiple comparisons when appropriate.

---

**Supplementary Stats Table: A compendium of all the statistical analyses associated with the figures.**

**Figure 1.**
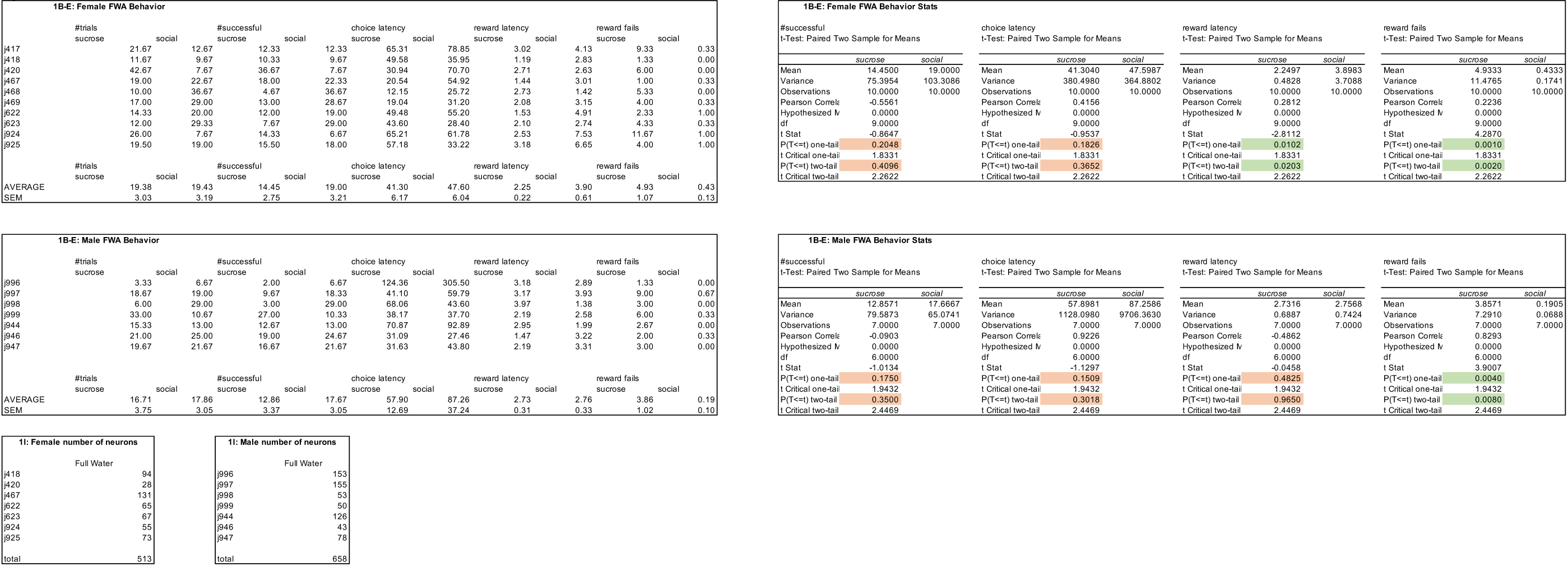

**Figure 2.**
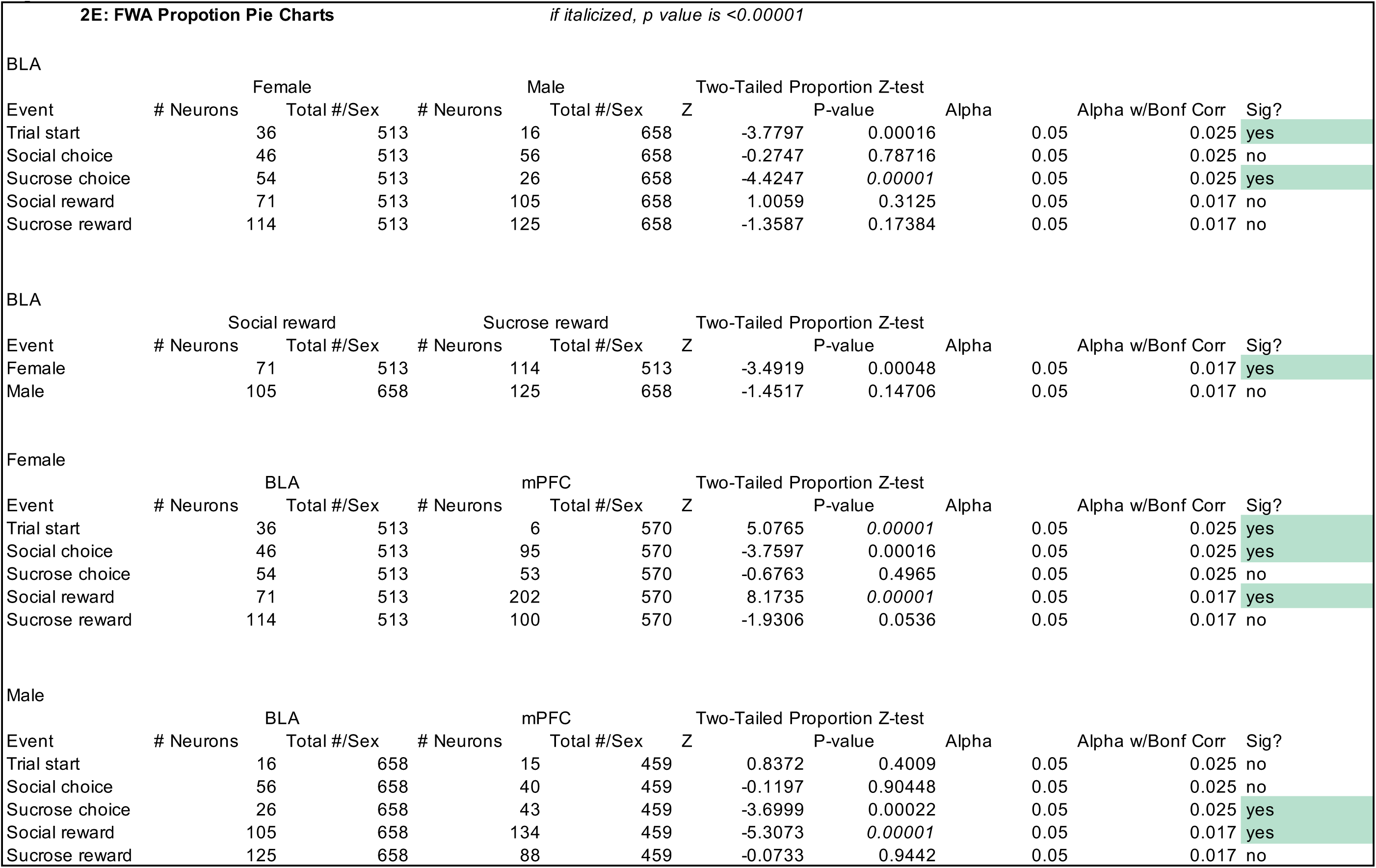

**Figure 3.**
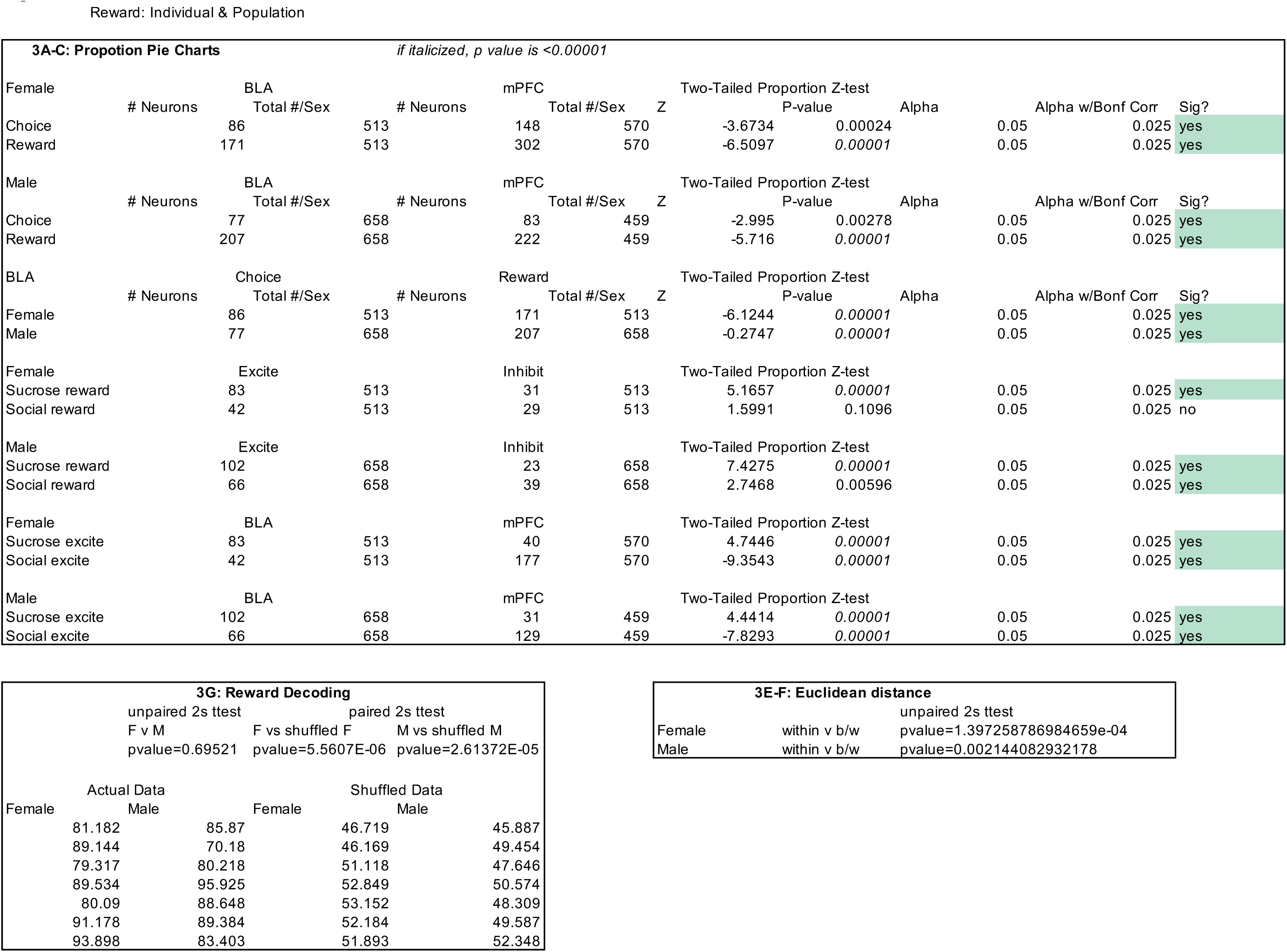

**Figure 4.**
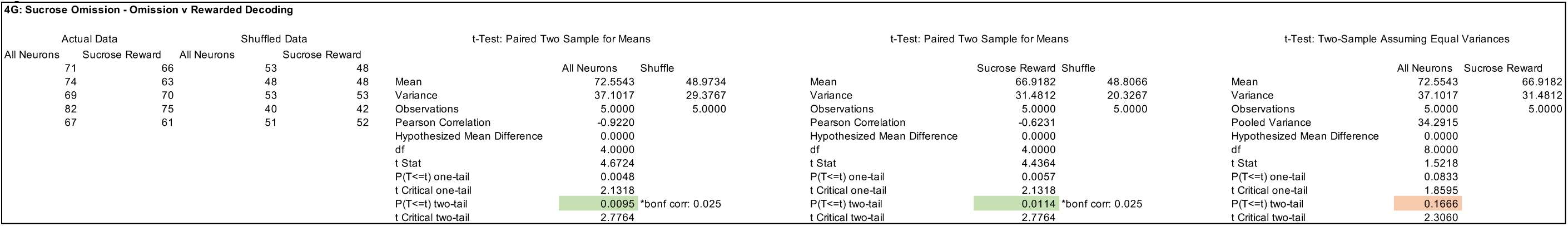

**Supplementary Figure 1.**
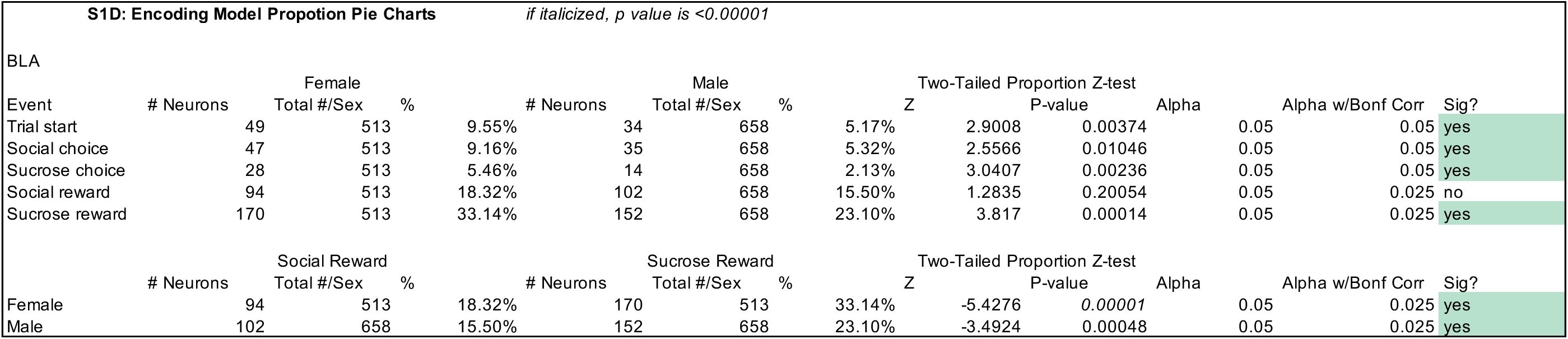

**Supplementary Figure 4.**
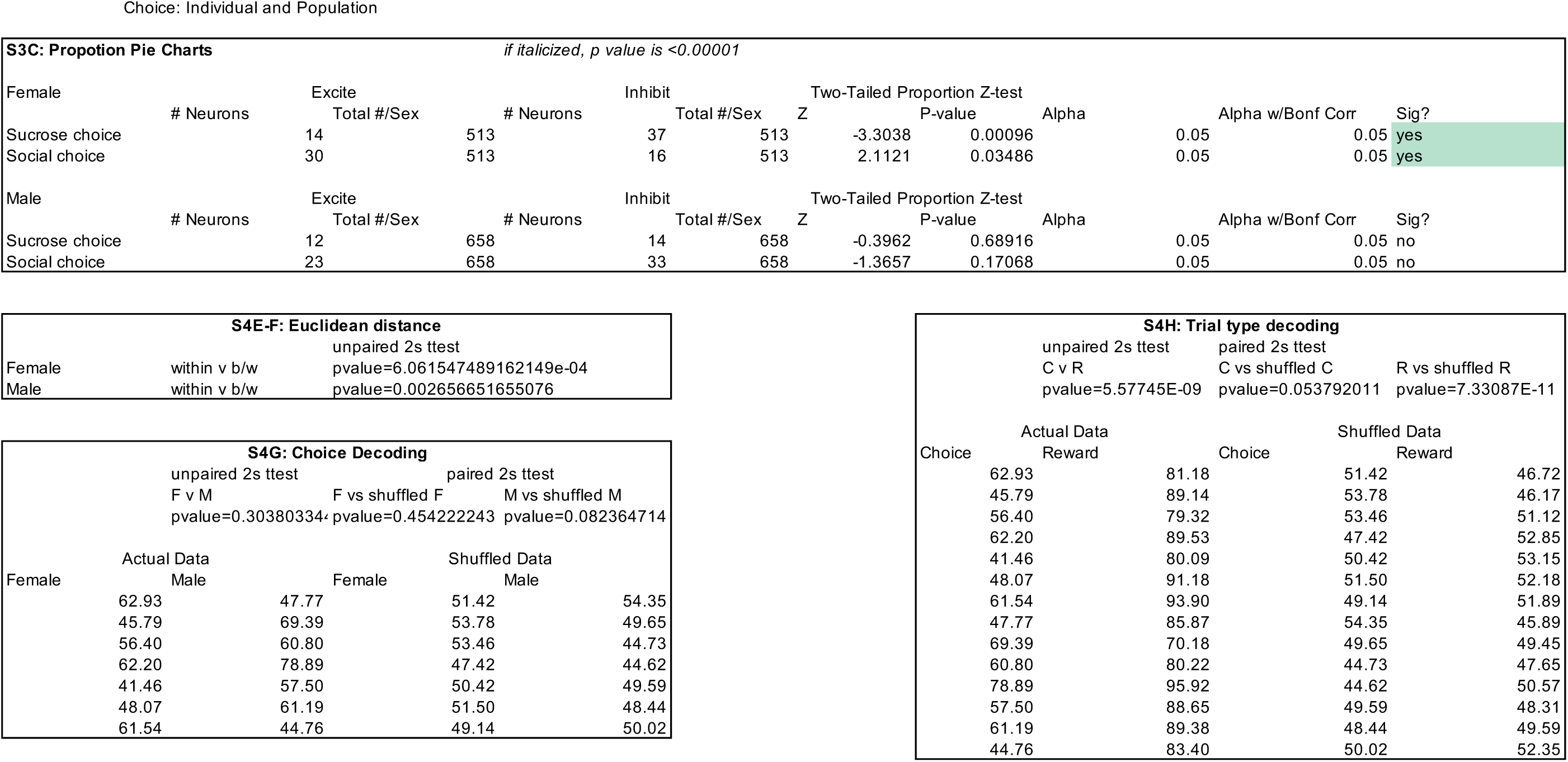

**Supplementary Figure 5.**
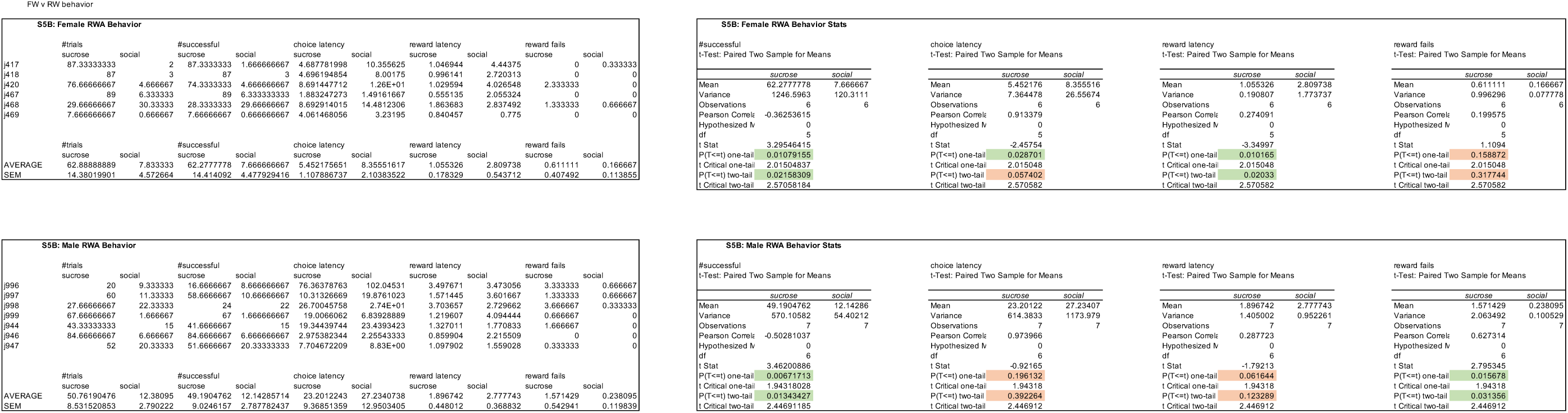

**Supplementary Figure 6.**
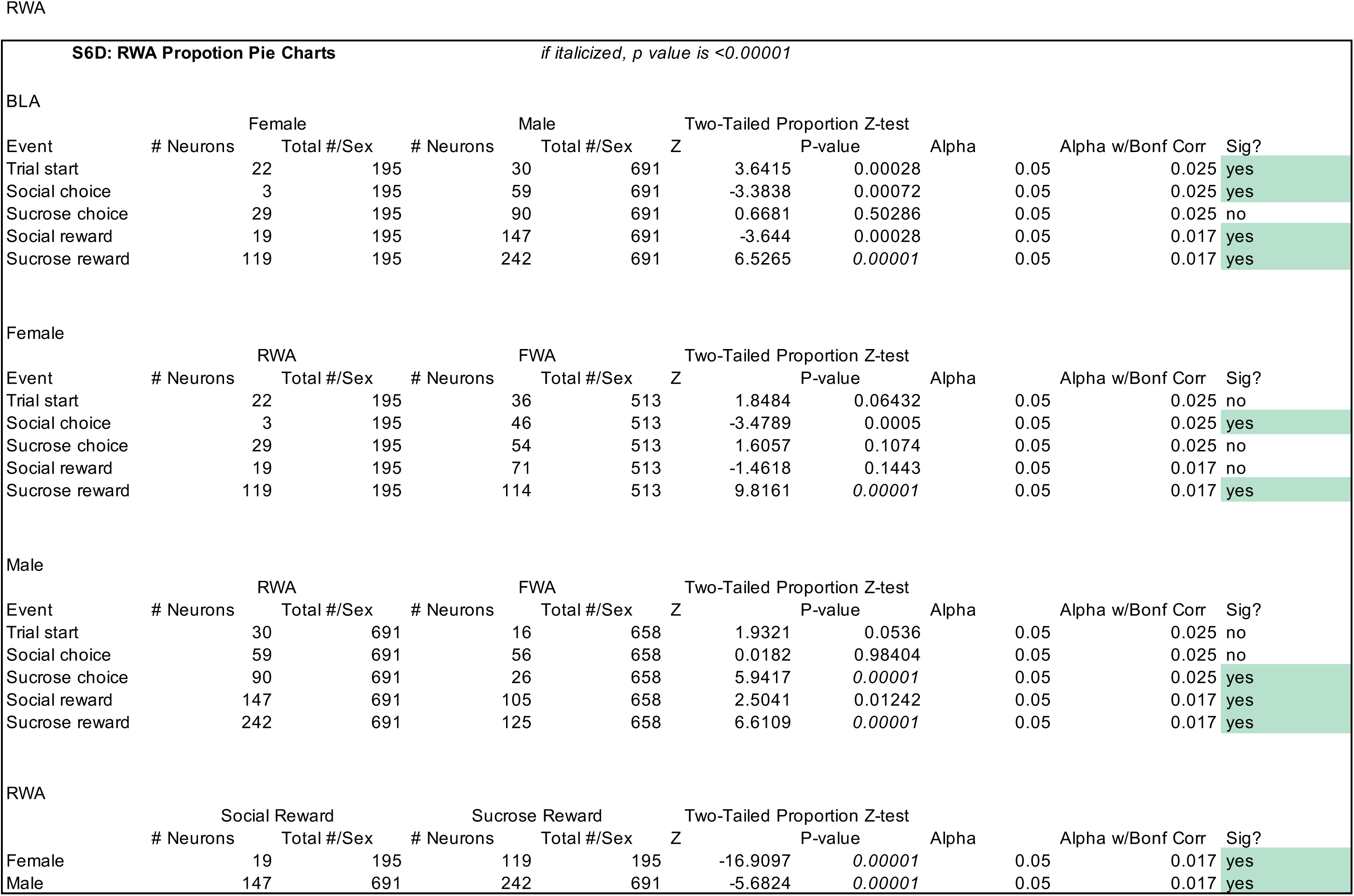

**Supplementary Figure 7.**
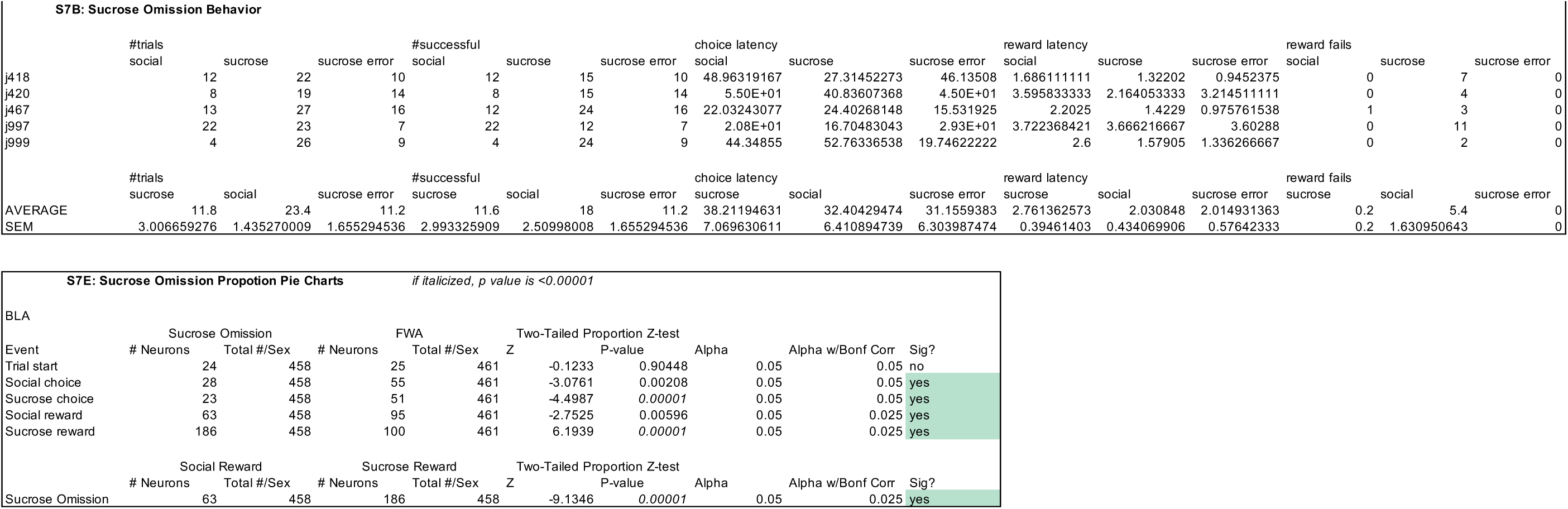

## References

1. Russo, S. J. & Nestler, E. J. The brain reward circuitry in mood disorders. Nature Reviews Neuroscience 14, 609–625 (2013).

2. Knowland, D. & Lim, B. K. Circuit-based frameworks of depressive behaviors: The role of reward circuitry and beyond. Pharmacology Biochemistry and Behavior 174, 42–52 (2018).

3. Yuan, Z. et al. A corticoamygdalar pathway controls reward devaluation and depression using dynamic inhibition code. Neuron 111, 3837–3853.e5 (2023).

4. Kumar, P. et al. Impaired reward prediction error encoding and striatal-midbrain connectivity in depression. Neuropsychopharmacology 43, 1581–1588 (2018).

5. Ng, T. H., Alloy, L. B. & Smith, D. V. Meta-analysis of reward processing in major depressive disorder reveals distinct abnormalities within the reward circuit. Translational Psychiatry 9, 1–10 (2019).

6. Bigot, M. et al. Disrupted basolateral amygdala circuits supports negative valence bias in depressive states. Translational Psychiatry 14, 382 (2024).

7. Dillon, D. G. & Pizzagalli, D. A. Mechanisms of Memory Disruption in Depression. Trends in neurosciences 41, 137 (2018).

8. Grueter, B. A., Rothwell, P. E. & Malenka, R. C. Integrating Synaptic Plasticity and Striatal Circuit Function in Addiction. Current Opinion in Neurobiology 22, 545 (2011).

9. Valence processing in the PFC: Reconciling circuit-level and systems-level views. In International Review of Neurobiology vol. 158 171–212 (Academic Press, 2021).

10. Konova, A. B. et al. Structural and behavioral correlates of abnormal encoding of money value in the sensorimotor striatum in cocaine addiction. European Journal of Neuroscience 36, 2979–2988 (2012).

11. Kalhan, S., Redish, A. D., Hester, R. & Garrido, M. I. A salience misattribution model for addictive-like behaviors. Neuroscience & Biobehavioral Reviews 125, 466–477 (2021).

12. Fraser, K. M., et al. Encoding and context-dependent control of reward consumption within the central nucleus of the amygdala. iScience 27, (2024).

13. Konova, A. B. et al. Reduced neural encoding of utility prediction errors in cocaine addiction. Neuron 111, (2023).

14. Lüscher, C. & Janak, P. H. Consolidating the Circuit Model for Addiction. Annual review of neuroscience 44, (2021).

15. Keiflin, R. & Janak, P. H. Dopamine Prediction Errors in Reward Learning and Addiction: From Theory to Neural Circuitry. Neuron 88, (2015).

16. Gold, J. M., Waltz, J. A., Prentice, K. J., Morris, S. E. & Heerey, E. A. Reward Processing in Schizophrenia: A Deficit in the Representation of Value. Schizophr Bull 34, 835–847 (2008).

17. Kirschner, M. et al. Deficits in context-dependent adaptive coding of reward in schizophrenia. npj Schizophrenia 2, 1–8 (2016).

18. Strauss, G. P., Waltz, J. A. & Gold, J. M. A Review of Reward Processing and Motivational Impairment in Schizophrenia. Schizophr Bull 40, S107–S116 (2013).

19. Vanes, L. D., Mouchlianitis, E., Collier, T., Averbeck, B. B. & Shergill, S. S. Differential neural reward mechanisms in treatment-responsive and treatment-resistant schizophrenia. Psychological Medicine 48, 2418–2427 (2018).

20. Lee, J., Jung, S., Park, I. & Kim, J.-J. Neural Basis of Anhedonia and Amotivation in Patients with Schizophrenia: The Role of Reward System. Current Neuropharmacology 13, 750–759 (2015).

21. Hernandez, A., Burton, A. C., O’Donnell, P., Schoenbaum, G. & Roesch, M. R. Altered Basolateral Amygdala Encoding in an Animal Model of Schizophrenia. J. Neurosci. 35, 6394–6400 (2015).

22. Zeng, J. et al. Neural substrates of reward anticipation and outcome in schizophrenia: a meta-analysis of fMRI findings in the monetary incentive delay task. Translational Psychiatry 12, 1–14 (2022).

23. Isaac, J. & Murugan, M. Interconnected neural circuits mediating social reward. Trends in neurosciences 47, (2024).

24. Lin, A., Adolphs, R. & Rangel, A. Social and monetary reward learning engage overlapping neural substrates. Soc Cogn Affect Neurosci 7, 274–281 (2011).

25. Isaac, J. et al. Sex differences in neural representations of social and nonsocial reward in the medial prefrontal cortex. Nat Commun 15, 8018 (2024).

26. Pinho, J. S., Cunliffe, V., Kareklas, K., Petri, G. & Oliveira, R. F. Social and asocial learning in zebrafish are encoded by a shared brain network that is differentially modulated by local activation. Communications Biology 6, 1–13 (2023).

27. O’Connell, L. A. & Hofmann, H. A. The Vertebrate mesolimbic reward system and social behavior network: A comparative synthesis. Journal of Comparative Neurology 519, 3599–3639 (2011).

28. Rocha-Almeida, F., Conde-Moro, A. R., Fernández-Ruiz, A., Delgado-García, J. M. & Gruart, A. Cortical and subcortical activities during food rewards versus social interaction in rats. Scientific Reports 15, 1–13 (2025).

29. Jennings, J. H. et al. Interacting neural ensembles in orbitofrontal cortex for social and feeding behaviour. Nature 565, (2019).

30. Vázquez, D., Schneider, K. N. & Roesch, M. R. Neural signals implicated in the processing of appetitive and aversive events in social and non-social contexts. Front. Syst. Neurosci. 16, 926388 (2022).

31. Willmore, L. et al. Overlapping representations of food and social stimuli in mouse VTA dopamine neurons. Neuron 111, 3541–3553.e8 (2023).

32. Isaac, J., Balasubramanian, H., Karkare, S. C., Schappaugh, N. & Murugan, M. Protocol for the quantitative assessment of social and nonsocial reward-seeking in mice using an automated two-choice operant assay. STAR Protoc 6, 103788 (2025).

33. Pignatelli, M. & Beyeler, A. Valence coding in amygdala circuits. Current Opinion in Behavioral Sciences 26, 97–106 (2019).

34. O’Neill, P.-K., Gore, F. & Salzman, C. D. Basolateral amygdala circuitry in positive and negative valence. Current Opinion in Neurobiology 49, 175–183 (2018).

35. Smith, D. M. & Torregrossa, M. M. Valence encoding in the amygdala influences motivated behavior. Behavioural Brain Research 411, 113370 (2021).

36. Fuster, J. M. & Uyeda, A. A. Reactivity of limbic neurons of the monkey to appetitive and aversive signals. Electroencephalography and Clinical Neurophysiology 30, 281–293 (1971).

37. Zhang, X. & Li, B. Population coding of valence in the basolateral amygdala. Nature Communications 9, 1–14 (2018).

38. Namburi, P., Al-Hasani, R., Calhoon, G. G., Bruchas, M. R. & Tye, K. M. Architectural Representation of Valence in the Limbic System. Neuropsychopharmacology 41, 1697–1715 (2015).

39. Pryce, C. R. Comparative evidence for the importance of the amygdala in regulating reward salience. Current Opinion in Behavioral Sciences 22, 76–81 (2018).

40. Janak, P. H. & Tye, K. M. From circuits to behaviour in the amygdala. Nature 517, 284–292 (2015).

41. Murray, E. A. The amygdala, reward and emotion. Trends in cognitive sciences 11, (2007).

42. Tian, Z. et al. The interhemispheric amygdala-accumbens circuit encodes negative valence in mice. Science (2024) doi:10.1126/science.adp7520.

43. Beyeler, A. et al. Organization of Valence-Encoding and Projection-Defined Neurons in the Basolateral Amygdala. Cell reports 22, 905 (2018).

44. Kyriazi, P., Headley, D. B. & Pare, D. Multi-dimensional Coding by Basolateral Amygdala Neurons. Neuron 99, 1315–1328.e5 (2018).

45. Kyriazi, P., Headley, D. B. & Paré, D. Different Multidimensional Representations across the Amygdalo-Prefrontal Network during an Approach-Avoidance Task. Neuron 107, 717 (2020).

46. Saez, A., Rigotti, M., Ostojic, S., Fusi, S. & Salzman, C. D. Abstract Context Representations in Primate Amygdala and Prefrontal Cortex. Neuron 87, (2015).

47. Piantadosi, S. C. et al. Holographic stimulation of opposing amygdala ensembles bidirectionally modulates valence-specific behavior via mutual inhibition. Neuron 112, (2024).

48. Shabel, S. J., Schairer, W., Donahue, R. J., Powell, V. & Janak, P. H. Similar neural activity during fear and disgust in the rat basolateral amygdala. PloS one 6, (2011).

49. Zelikowsky, M., Hersman, S., Chawla, M. K., Barnes, C. A. & Fanselow, M. S. Neuronal ensembles in amygdala, hippocampus, and prefrontal cortex track differential components of contextual fear. The Journal of neuroscience : the official journal of the Society for Neuroscience 34, (2014).

50. Brockett, A. T., Vázquez, D. & Roesch, M. R. Prediction errors and valence: From single units to multidimensional encoding in the amygdala. Behavioural brain research 404, (2021).

51. Paton, J. J., Belova, M. A., Morrison, S. E. & Salzman, C. D. The primate amygdala represents the positive and negative value of visual stimuli during learning. Nature 439, (2006).

52. Shabel, S. J. & Janak, P. H. Substantial similarity in amygdala neuronal activity during conditioned appetitive and aversive emotional arousal. Proceedings of the National Academy of Sciences of the United States of America 106, (2009).

53. Fustiñana, M. S., Eichlisberger, T., Bouwmeester, T., Bitterman, Y. & Lüthi, A. State-dependent encoding of exploratory behaviour in the amygdala. Nature 592, 267–271 (2021).

54. Kietzman, H. W., Trinoskey-Rice, G., Seo, E. H., Guo, J. & Gourley, S. L. Neuronal Ensembles in the Amygdala Allow Social Information to Motivate Later Decisions. J. Neurosci. 44, (2024).

55. Morris, J. S. et al. A differential neural response in the human amygdala to fearful and happy facial expressions. Nature 383, (1996).

56. Felix-Ortiz, A. C. & Tye, K. M. Amygdala Inputs to the Ventral Hippocampus Bidirectionally Modulate Social Behavior. J. Neurosci. 34, 586–595 (2014).

57. Taylor, S. L., Stanek, L. M., Ressler, K. J. & Huhman, K. L. Differential brain-derived neurotrophic factor expression in limbic brain regions following social defeat or territorial aggression. Behavioral neuroscience 125, (2011).

58. Li, M. et al. Ketamine ameliorates post-traumatic social avoidance by erasing the traumatic memory encoded in VTA-innervated BLA engram cells. Neuron 112, (2024).

59. Terranova, J. I. et al. Hippocampal-amygdala memory circuits govern experience-dependent observational fear. Neuron 110, (2022).

60. Haruno, M. & Frith, C. D. Activity in the amygdala elicited by unfair divisions predicts social value orientation. Nature neuroscience 13, (2010).

61. Gadziola, M. A., Shanbhag, S. J. & Wenstrup, J. J. Two distinct representations of social vocalizations in the basolateral amygdala. Journal of Neurophysiology (2016) doi:10.1152/jn.00953.2015.

62. Belova, M. A., Paton, J. J. & Daniel Salzman, C. Moment-to-Moment Tracking of State Value in the Amygdala. J. Neurosci. 28, 10023–10030 (2008).

63. Schoenbaum, G., Chiba, A. A. & Gallagher, M. Orbitofrontal cortex and basolateral amygdala encode expected outcomes during learning. Nature Neuroscience 1, 155–159 (1998).

64. Kim, J., Pignatelli, M., Xu, S., Itohara, S. & Tonegawa, S. Antagonistic negative and positive neurons of the basolateral amygdala. Nature Neuroscience 19, 1636–1646 (2016).

65. Rogan, M. T., Stäubli, U. V. & LeDoux, J. E. Fear conditioning induces associative long-term potentiation in the amygdala. Nature 390, (1997).

66. Parsana, A. J., Li, N. & Brown, T. H. Positive and Negative Ultrasonic Social Signals Elicit Opposing Firing Patterns in Rat Amygdala. Behavioural brain research 226, 77 (2011).

67. Wassum, K. M. & Izquierdo, A. The basolateral amygdala in reward learning and addiction. Neuroscience & Biobehavioral Reviews 57, 271–283 (2015).

68. Wassum, K. M. Amygdala-cortical collaboration in reward learning and decision making. (2022) doi:10.7554/eLife.80926.

69. Beyeler, A. et al. Divergent Routing of Positive and Negative Information from the Amygdala during Memory Retrieval. Neuron 90, (2016).

70. Amaya, K. A., Teboul, E., Weiss, G. L., Antonoudiou, P. & Maguire, J. L. Basolateral amygdala parvalbumin interneurons coordinate oscillations to drive reward behaviors. Current biology : CB 34, (2024).

71. Gore, F. et al. Neural Representations of Unconditioned Stimuli in Basolateral Amygdala Mediate Innate and Learned Responses. Cell 162, 134–145 (2015).

72. Hinz, J. et al. Stimulus-specific and adaptive value representations in the basolateral amygdala in male mice. Nature Communications 16, 1–12 (2025).

73. Levy, D. J. & Glimcher, P. W. The root of all value: a neural common currency for choice. Current opinion in neurobiology 22, (2012).

74. Tye, K. M. et al. Mixed selectivity: Cellular computations for complexity. Neuron 112, 2289–2303 (2024).

75. Gothard, K. M. Multidimensional processing in the amygdala. Nature Reviews Neuroscience 21, 565–575 (2020).

76. Maren, S. Parsing Reward and Aversion in the Amygdala. Neuron 90, (2016).

77. Tye, K. M. Neural Circuit Motifs in Valence Processing. Neuron 100, 436–452 (2018).

78. Munuera, J., Rigotti, M. & Salzman, C. D. Shared neural coding for social hierarchy and reward value in primate amygdala. Nat Neurosci 21, 415–423 (2018).

79. Mazuski, C. & O’Keefe, J. Representation of ethological events by basolateral amygdala neurons. Cell reports 39, (2022).

80. McGarry, L. M. & Carter, A. G. Prefrontal Cortex Drives Distinct Projection Neurons in the Basolateral Amygdala. Cell reports 21, (2017).

81. Yizhar, O. & Klavir, O. Reciprocal amygdala-prefrontal interactions in learning. Current opinion in neurobiology 52, (2018).

82. Fetterly, T. L., Catalfio, A. M. & Ferrario, C. R. Effects of junk-food on food-motivated behavior and nucleus accumbens glutamate plasticity; insights into the mechanism of calcium-permeable AMPA receptor recruitment. Neuropharmacology 242, 109772 (2024).

83. Moorman, D. E. & Aston-Jones, G. Prefrontal neurons encode context-based response execution and inhibition in reward seeking and extinction. Proceedings of the National Academy of Sciences 112, 9472–9477 (2015).

84. Murugan, M. et al. Combined Social and Spatial Coding in a Descending Projection from the Prefrontal Cortex. Cell 171, (2017).

85. Kingsbury, L. et al. Correlated Neural Activity and Encoding of Behavior across Brains of Socially Interacting Animals. Cell 178, (2019).

86. Zhang, W. & Yartsev, M. M. Correlated Neural Activity across the Brains of Socially Interacting Bats. Cell 178, (2019).

87. Jabarin, R., Mohapatra, A. N., Ray, N., Netser, S. & Wagner, S. Distinct prelimbic cortex neuronal responses to emotional states of others drive emotion recognition in adult mice. Current biology : CB 35, (2025).

88. Amadei, E. A. et al. Dynamic corticostriatal activity biases social bonding in monogamous female prairie voles. Nature 546, 297–301 (2017).

89. Ambroggi, F., Ishikawa, A., Fields, H. L. & Nicola, S. M. Basolateral amygdala neurons facilitate reward-seeking behavior by exciting nucleus accumbens neurons. Neuron 59, 648 (2008).

90. Ishikawa, A., Ambroggi, F., Nicola, S. M. & Fields, H. L. Contributions of the amygdala and medial prefrontal cortex to incentive cue responding. Neuroscience 155, 573 (2008).

91. Stefanik, M. T. & Kalivas, P. W. Optogenetic dissection of basolateral amygdala projections during cue-induced reinstatement of cocaine seeking. Front. Behav. Neurosci. 7, 74455 (2013).

92. Mora, F. & Myers, R. D. Brain self-stimulation: direct evidence for the involvement of dopamine in the prefrontal cortex. Science 197, 1387–1389 (1977).

93. You, Z. B., Tzschentke, T. M., Brodin, E. & Wise, R. A. Electrical stimulation of the prefrontal cortex increases cholecystokinin, glutamate, and dopamine release in the nucleus accumbens: an in vivo microdialysis study in freely moving rats. J. Neurosci. 18, 6492–6500 (1998).

94. Zhang, Y.-Q. et al. Abused drug-induced intracranial self-stimulation is correlated with the alteration of dopamine transporter availability in the medial prefrontal cortex and nucleus accumbens of mice. Biomedicine & Pharmacotherapy 169, 115860 (2023).

95. Britt, J. P. et al. Synaptic and behavioral profile of multiple glutamatergic inputs to the nucleus accumbens. Neuron 76, (2012).

96. Stuber, G. D. et al. Amygdala to nucleus accumbens excitatory transmission facilitates reward seeking. Nature 475, 377 (2011).

97. Zhou, K. et al. Reward and aversion processing by input-defined parallel nucleus accumbens circuits in mice. Nature Communications 13, 1–12 (2022).

98. He, Y., Huang, Y. H., Schlüter, O. M. & Dong, Y. Cue- versus reward-encoding basolateral amygdala projections to nucleus accumbens. (2023) doi:10.7554/eLife.89766.

99. Liu, Y. et al. A molecularly defined mPFC-BLA circuit specifically regulates social novelty preference. Science Advances (2025) doi:10.1126/sciadv.adt9008.

100. Huang, W. C., Zucca, A., Levy, J. & Page, D. T. Social Behavior Is Modulated by Valence-Encoding mPFC-Amygdala Sub-circuitry. Cell reports 32, (2020).

101. Huang, W. C., Chen, Y. & Page, D. T. Hyperconnectivity of prefrontal cortex to amygdala projections in a mouse model of macrocephaly/autism syndrome. Nature communications 7, (2016).

102. Tan, Y. et al. Oxytocin Receptors Are Expressed by Glutamatergic Prefrontal Cortical Neurons That Selectively Modulate Social Recognition. The Journal of neuroscience : the official journal of the Society for Neuroscience 39, (2019).

103. Felix-Ortiz, A. C., Burgos-Robles, A., Bhagat, N. D., Leppla, C. A. & Tye, K. M. Bidirectional modulation of anxiety-related and social behaviors by amygdala projections to the medial prefrontal cortex. Neuroscience 321, (2016).

104. Grossman, Y. S. et al. Structure and function differences in the prelimbic cortex to basolateral amygdala circuit mediate trait vulnerability in a novel model of acute social defeat stress in male mice. Neuropsychopharmacology 47, 788–799 (2021).

105. Kumaran, D., Banino, A., Blundell, C., Hassabis, D. & Dayan, P. Computations Underlying Social Hierarchy Learning: Distinct Neural Mechanisms for Updating and Representing Self-Relevant Information. Neuron 92, (2016).

106. Diaz, V. & Lin, D. Neural circuits for coping with social defeat. Current opinion in neurobiology 60, 99 (2019).

107. Xin, Q. et al. Deconstructing the neural circuit underlying social hierarchy in mice. Neuron 113, (2025).

108. Keefer, S. E. & Petrovich, G. D. Distinct recruitment of basolateral amygdala-medial prefrontal cortex pathways across Pavlovian appetitive conditioning. Neurobiology of learning and memory 141, (2017).

109. Lee, J. et al. Neural dynamics of emergent social roles in collective foraging by mice. (2025) doi:10.1101/2025.04.24.650547.

110. Keefer, S. E. & Petrovich, G. D. The basolateral amygdala-medial prefrontal cortex circuitry regulates behavioral flexibility during appetitive reversal learning. Behavioral neuroscience 134, (2020).

111. Petrovich, G. D. & Gallagher, M. Control of food consumption by learned cues: a forebrain-hypothalamic network. Physiology & behavior 91, (2007).

112. Reppucci, C. J. & Petrovich, G. D. Organization of connections between the amygdala, medial prefrontal cortex, and lateral hypothalamus: a single and double retrograde tracing study in rats. Brain Structure and Function 221, 2937–2962 (2015).

113. Kietzman, H. W., Trinoskey-Rice, G., Blumenthal, S. A., Guo, J. D. & Gourley, S. L. Social incentivization of instrumental choice in mice requires amygdala-prelimbic cortex-nucleus accumbens connectivity. Nature Communications 13, 1–11 (2022).

114. Burgos-Robles, A. et al. Amygdala inputs to prefrontal cortex guide behavior amid conflicting cues of reward and punishment. Nature Neuroscience 20, 824–835 (2017).

115. Liu, N., Li, Y. F., Zhao, X. T., Li, Y. H. & Cui, R. S. Inhibition of the basolateral amygdala to prelimbic cortex pathway enhances risk-taking during risky decision-making shock task in rats. Physiology & behavior 292, (2025).

116. Scheggia, D. et al. Reciprocal cortico-amygdala connections regulate prosocial and selfish choices in mice. Nature Neuroscience 25, 1505–1518 (2022).

117. Chefer, V. I., Wang, R. & Shippenberg, T. S. Basolateral Amygdala-Driven Augmentation of Medial Prefrontal Cortex GABAergic Neurotransmission in Response to Environmental Stimuli Associated with Cocaine Administration. Neuropsychopharmacology 36, 2018–2029 (2011).

118. Patel, R. R. et al. Social isolation recruits amygdala-cortical circuitry to escalate alcohol drinking. Research square (2024) doi:10.21203/rs.3.rs-4033115/v1.

119. Rosenberger, L. A. et al. The Human Basolateral Amygdala Is Indispensable for Social Experiential Learning. Current Biology 29, 3532–3537.e3 (2019).

120. Rodriguez, L. A. et al. The basolateral amygdala to lateral septum circuit is critical for regulating social novelty in mice. Neuropsychopharmacology 48, 529–539 (2022).

121. Namburi, P. et al. A circuit mechanism for differentiating positive and negative associations. Nature 520, 675–678 (2015).

122. LeDoux, J. The Emotional Brain, Fear, and the Amygdala. Cellular and Molecular Neurobiology 23, 727–738 (2003).

123. Averbeck, B. B. & Costa, V. D. Motivational neural circuits underlying reinforcement learning. Nature Neuroscience 20, 505–512 (2017).

124. Servonnet, A., Hernandez, G., El Hage, C., Rompré, P. P. & Samaha, A. N. Optogenetic Activation of the Basolateral Amygdala Promotes Both Appetitive Conditioning and the Instrumental Pursuit of Reward Cues. The Journal of neuroscience : the official journal of the Society for Neuroscience 40, (2020).

125. Lim, H. et al. Genetically- and spatially-defined basolateral amygdala neurons control food consumption and social interaction. Nature Communications 15, 1–22 (2024).

126. Liu, J. et al. Neural Coding of Appetitive Food Experiences in the Amygdala. Neurobiol Learn Mem 155, 261–275 (2018).

127. Zhang, X., Kim, J. & Tonegawa, S. Amygdala Reward Neurons Form and Store Fear Extinction Memory. Neuron 105, 1077–1093.e7 (2020).

128. Adolphs, R., Russell, J. A. & Tranel, D. A Role for the Human Amygdala in Recognizing Emotional Arousal From Unpleasant Stimuli. Psychological Science (1999) doi:10.1111/1467-9280.00126.

129. Zald, D. H. & Pardo, J. V. Emotion, olfaction, and the human amygdala: Amygdala activation during aversive olfactory stimulation. Proceedings of the National Academy of Sciences 94, 4119–4124 (1997).

130. Davis, M. The role of the amygdala in fear and anxiety. Annual review of neuroscience 15, (1992).

131. Blanchard, D. C. & Blanchard, R. J. Innate and conditioned reactions to threat in rats with amygdaloid lesions. Journal of comparative and physiological psychology 81, (1972).

132. Nachman, M. & Ashe, J. H. Effects of basolateral amygdala lesions on neophobia, learned taste aversions, and sodium appetite in rats. J Comp Physiol Psychol 87, 622–643 (1974).

133. Huff, M. L., Emmons, E. B., Narayanan, N. S. & LaLumiere, R. T. Basolateral amygdala projections to ventral hippocampus modulate the consolidation of footshock, but not contextual, learning in rats. Learn. Mem. 23, 51–60 (2016).

134. Sierra-Mercado, D., Padilla-Coreano, N. & Quirk, G. J. Dissociable Roles of Prelimbic and Infralimbic Cortices, Ventral Hippocampus, and Basolateral Amygdala in the Expression and Extinction of Conditioned Fear. Neuropsychopharmacology 36, 529–538 (2010).

135. Gangopadhyay, P., Chawla, M., Dal Monte, O. & Chang, S. W. C. Prefrontal–amygdala circuits in social decision-making. Nature Neuroscience 24, 5–18 (2020).

136. Li, M., Jiang, Y. Q. & Sun, Q. A circuit from the basolateral amygdala to hippocampal CA3 regulates social behavior. Current biology : CB (2025) doi:10.1016/j.cub.2025.07.059.

137. Chang, S. W. C. et al. Neural mechanisms of social decision-making in the primate amygdala. Proceedings of the National Academy of Sciences 112, 16012–16017 (2015).

138. Nerio-Morales, L. K., Boender, A. J., Young, L. J., Lamprea, M. R. & Smith, A. S. Limbic oxytocin receptor expression alters molecular signaling and social avoidance behavior in female prairie voles (Microtus ochrogaster). Front. Neurosci. 18, 1409316 (2024).

139. Ferrara, N. C. & Opendak, M. Amygdala circuit transitions supporting developmentally-appropriate social behavior. Neurobiol Learn Mem 201, 107762 (2023).

140. Allsop, S. A. et al. Corticoamygdala transfer of socially-derived information gates observational learning. Cell 173, 1329 (2018).

141. Warthen, K. G. et al. Sex differences in the human reward system: convergent behavioral, autonomic and neural evidence. Soc Cogn Affect Neurosci 15, 789–801 (2020).

142. Vantrease, J. E. et al. Sex Differences in the Activity of Basolateral Amygdalar Neurons That Project to the Bed Nucleus of the Stria Terminalis and Their Role in Anticipatory Anxiety. The Journal of neuroscience : the official journal of the Society for Neuroscience 42, (2022).

143. Guily, P., Lassalle, O., Chavis, P. & Manzoni, O. J. Sex-specific divergent maturational trajectories in the postnatal rat basolateral amygdala. iScience 25, (2022).

144. Manion, M. T. C., Glasper, E. R. & Wang, K. H. A sex difference in mouse dopaminergic projections from the midbrain to basolateral amygdala. Biology of Sex Differences 13, 1–13 (2022).

145. Taniguchi, L., et al. Sex-and Stress-Dependent Plasticity of a Corticotropin Releasing Hormone / GABA Projection from the Basolateral Amygdala to Nucleus Accumbens that Mediates Reward Behaviors. bioRxiv : the preprint server for biology (2024) doi:10.1101/2024.11.30.626183.

146. Song, Z., Swarna, S. & Manns, J. R. Prioritization of social information by the basolateral amygdala in rats. Neurobiology of learning and memory 184, 107489 (2021).

147. Opendak, M. et al. Bidirectional control of infant rat social behavior via dopaminergic innervation of the basolateral amygdala. Neuron 109, 4018–4035.e7 (2021).

148. Kim, J., Zhang, X., Muralidhar, S., LeBlanc, S. A. & Tonegawa, S. Basolateral to Central Amygdala Neural Circuits for Appetitive Behaviors. Neuron 93, (2017).

149. Khalil, V. et al. Subcortico-amygdala pathway processes innate and learned threats. (2023) doi:10.7554/eLife.85459.

150. Jones, J. L. et al. Basolateral amygdala modulates terminal dopamine release in the nucleus accumbens and conditioned responding. Biological psychiatry 67, (2010).

151. Sun, L. et al. Reactivating a positive feedback loop VTA-BLA-NAc circuit associated with positive experience ameliorates the attenuated reward sensitivity induced by chronic stress. Neurobiology of stress 15, (2021).

152. Morse, A. K. et al. Basolateral Amygdala Drives a GPCR-Mediated Striatal Memory Necessary for Predictive Learning to Influence Choice. Neuron 106, (2020).

153. Laviolette, S. R. & Grace, A. A. The roles of cannabinoid and dopamine receptor systems in neural emotional learning circuits: implications for schizophrenia and addiction. Cellular and molecular life sciences : CMLS 63, (2006).

154. Rosen, L. G., Sun, N., Rushlow, W. & Laviolette, S. R. Molecular and neuronal plasticity mechanisms in the amygdala-prefrontal cortical circuit: implications for opiate addiction memory formation. Front. Neurosci. 9, 164878 (2015).

155. Morel, C. et al. Midbrain projection to the basolateral amygdala encodes anxiety-like but not depression-like behaviors. Nature Communications 13, 1–13 (2022).

156. Sias, A. C. et al. Dopamine projections to the basolateral amygdala drive the encoding of identity-specific reward memories. Nature neuroscience 27, (2024).

157. Pujara, M. S., Ciesinski, N. K., Reyelts, J. F., Rhodes, S. E. V. & Murray, E. A. Selective Prefrontal-Amygdala Circuit Interactions Underlie Social and Nonsocial Valuation in Rhesus Macaques. The Journal of neuroscience : the official journal of the Society for Neuroscience 42, (2022).

158. Laviolette, S. R., Lipski, W. J. & Grace, A. A. A subpopulation of neurons in the medial prefrontal cortex encodes emotional learning with burst and frequency codes through a dopamine D4 receptor-dependent basolateral amygdala input. The Journal of neuroscience : the official journal of the Society for Neuroscience 25, (2005).

159. Rosenkranz, J. A. & Grace, A. A. Cellular mechanisms of infralimbic and prelimbic prefrontal cortical inhibition and dopaminergic modulation of basolateral amygdala neurons in vivo. The Journal of neuroscience : the official journal of the Society for Neuroscience 22, (2002).

160. Wei, C. A. O., Huiyi, L. I. & Jianhong, L. U. O. Prefrontal cortical circuits in social behaviors: an overview. Journal of Zhejiang University. Science. B 25, 941 (2024).

161. Grossmann, T. Novel Insights into the Social Functions of the Medial Prefrontal Cortex during Infancy. eNeuro 12, ENEURO.0458–24.2025 (2025).

162. Ko, J. Neuroanatomical Substrates of Rodent Social Behavior: The Medial Prefrontal Cortex and Its Projection Patterns. Front. Neural Circuits 11, 276485 (2017).

163. Levy, D. R. et al. Dynamics of social representation in the mouse prefrontal cortex. Nature Neuroscience 22, 2013–2022 (2019).

164. Pastor, V. & Medina, J. H. Medial prefrontal cortical control of reward- and aversion-based behavioral output: Bottom-up modulation. The European journal of neuroscience 53, (2021).

165. Price, J. L. Definition of the orbital cortex in relation to specific connections with limbic and visceral structures and other cortical regions. Annals of the New York Academy of Sciences 1121, (2007).

166. Pavuluri, M., Volpe, K. & Yuen, A. Nucleus Accumbens and Its Role in Reward and Emotional Circuitry: A Potential Hot Mess in Substance Use and Emotional Disorders. AIMS Neuroscience 4, 52–70 (2017).

167. Mannella, F., Gurney, K. & Baldassarre, G. The nucleus accumbens as a nexus between values and goals in goal-directed behavior: a review and a new hypothesis. Front. Behav. Neurosci. 7, 51817 (2013).

168. Gruber, A. J. & McDonald, R. J. Context, emotion, and the strategic pursuit of goals: interactions among multiple brain systems controlling motivated behavior. Frontiers in behavioral neuroscience 6, (2012).

169. Scofield, M. D. et al. The Nucleus Accumbens: Mechanisms of Addiction across Drug Classes Reflect the Importance of Glutamate Homeostasis. Pharmacological Reviews 68, 816 (2016).

170. Zhao, P., et al. Accelerated social representational drift in the nucleus accumbens in a model of autism. bioRxiv : the preprint server for biology (2023) doi:10.1101/2023.08.05.552133.

171. Chen, C. S. et al. Divergent Strategies for Learning in Males and Females. Current biology : CB 31, (2021).

172. Cox, J. et al. A neural substrate of sex-dependent modulation of motivation. Nat Neurosci 26, 274–284 (2023).

173. Vahaba, D. M., Halstead, E. R., Donaldson, Z. R., Ahern, T. H. & Beery, A. K. Sex differences in the reward value of familiar mates in prairie voles. Genes, brain, and behavior 21, (2022).

174. Cognuck, S. Q. et al. Sex- and age-dependent differences in the hormone and drinking responses to water deprivation. American Journal of Physiology-Regulatory, Integrative and Comparative Physiology (2020) doi:10.1152/ajpregu.00303.2019.

175. Kaufman, S. A comparison of the dipsogenic responses of male and female rats to a variety of stimuli. Canadian Journal of Physiology and Pharmacology (2011) doi:10.1139/y80-179.

176. Hintiryan, H. et al. Connectivity characterization of the mouse basolateral amygdalar complex. Nature Communications 12, 1–25 (2021).

177. Pitkänen, A., Savander, V. & LeDoux, J. E. Organization of intra-amygdaloid circuitries in the rat: an emerging framework for understanding functions of the amygdala. Trends in neurosciences 20, (1997).

178. Hochgerner, H. et al. Neuronal types in the mouse amygdala and their transcriptional response to fear conditioning. Nature Neuroscience 26, 2237–2249 (2023).

179. Yang, Y. & Wang, J.-Z. From Structure to Behavior in Basolateral Amygdala-Hippocampus Circuits. Front. Neural Circuits 11, 281732 (2017).

180. Rodrigues, S. M., Schafe, G. E. & LeDoux, J. E. Molecular mechanisms underlying emotional learning and memory in the lateral amygdala. Neuron 44, (2004).

181. Spampanato, J., Polepalli, J. & Sah, P. Interneurons in the basolateral amygdala. Neuropharmacology 60, (2011).

182. Polepalli, J. S., Gooch, H. & Sah, P. Diversity of interneurons in the lateral and basal amygdala. npj Science of Learning 5, 1–9 (2020).

183. Rainnie, D. G., Asprodini, E. K. & Shinnick-Gallagher, P. Excitatory transmission in the basolateral amygdala. Journal of Neurophysiology (1991) doi:10.1152/jn.1991.66.3.986.

184. McDonald, A. J. Functional neuroanatomy of basal forebrain projections to the basolateral amygdala: Transmitters, receptors, and neuronal subpopulations. Journal of Neuroscience Research 102, e25318 (2024).

185. Carlsen, J. New perspectives on the functional anatomical organization of the basolateral amygdala. Acta Neurol Scand Suppl 122, 1–27 (1989).

186. Lanuza, E., Belekhova, M., Martínez-Marcos, A., Font, C. & Martínez-García, F. Identification of the reptilian basolateral amygdala: an anatomical investigation of the afferents to the posterior dorsal ventricular ridge of the lizard Podarcis hispanica. The European journal of neuroscience 10, (1998).

187. Sniffen, S. E. et al. Directing negative emotional states through parallel genetically-distinct basolateral amygdala pathways to ventral striatum subregions. Molecular Psychiatry 1–14 (2025).

188. Nakashima, M., Ikegaya, Y. & Morikawa, S. Genetic labeling of axo-axonic cells in the basolateral amygdala. Neuroscience research 178, (2022).

189. Jhaveri, D. J. et al. Evidence for newly generated interneurons in the basolateral amygdala of adult mice. Molecular Psychiatry 23, 521–532 (2017).

190. Morikawa, S., Katori, K., Takeuchi, H. & Ikegaya, Y. Brain-wide mapping of presynaptic inputs to basolateral amygdala neurons. The Journal of comparative neurology 529, (2021).

191. Ping, A. et al. Brainwide mesoscale functional networks revealed by focal infrared neural stimulation of the amygdala. National science review 12, (2024).

192. O’Leary, T. P. et al. Extensive and spatially variable within-cell-type heterogeneity across the basolateral amygdala. (2020) doi:10.7554/eLife.59003.

193. Power, J. M. & Sah, P. Dendritic spine heterogeneity and calcium dynamics in basolateral amygdala principal neurons. Journal of Neurophysiology (2014) doi:10.1152/jn.00770.2013.

194. McDonald, A. J. Neuronal organization of the lateral and basolateral amygdaloid nuclei in the rat. The Journal of comparative neurology 222, (1984).

195. Pitkänen, A., Jolkkonen, E. & Kemppainen, S. Anatomic heterogeneity of the rat amygdaloid complex. Folia Morphologica 59, 1–24 (2000).

196. Butler, R. K., Oliver, E. M., Fadel, J. R. & Wilson, M. A. Hemispheric differences in the number of parvalbumin-positive neurons in subdivisions of the rat basolateral amygdala complex. Brain Research 1678, 214–219 (2018).

197. Totty, M. S. et al. Transcriptomic diversity of amygdalar subdivisions across humans and nonhuman primates. bioRxiv 2024.10.18.618721 (2024) doi:10.1101/2024.10.18.618721.

198. Felix-Ortiz, A. C. et al. BLA to vHPC inputs modulate anxiety-related behaviors. Neuron 79, (2013).

199. Prager, E. M., Bergstrom, H. C., Wynn, G. H. & Braga, M. F. M. The Basolateral Amygdala GABAergic System in Health and Disease. Journal of neuroscience research 94, 548 (2015).

200. Hájos, N. Interneuron Types and Their Circuits in the Basolateral Amygdala. Frontiers in Neural Circuits 15, 687257 (2021).

201. Asim, M., Wang, H., Waris, A. & He, J. Basolateral amygdala parvalbumin and cholecystokinin-expressing GABAergic neurons modulate depressive and anxiety-like behaviors. Translational Psychiatry 14, 1–13 (2024).

202. Lüthi, A. & Lüscher, C. Pathological circuit function underlying addiction and anxiety disorders. Nature Neuroscience 17, 1635–1643 (2014).

203. Tuna, T. et al. Basal forebrain innervation of the amygdala: an anatomical and computational exploration. Brain Structure and Function 230, 1–30 (2025).

204. Antonoudiou, P. et al. Biased Information Routing Through the Basolateral Amygdala, Altered Valence Processing, and Impaired Affective States Associated With Psychiatric Illnesses. Biological Psychiatry 97, 764–774 (2025).

205. Malvaez, M., Shieh, C., Murphy, M. D., Greenfield, V. Y. & Wassum, K. M. Distinct cortical– amygdala projections drive reward value encoding and retrieval. Nature Neuroscience 22, 762–769 (2019).

206. McGarry, L. M. & Carter, A. G. Inhibitory Gating of Basolateral Amygdala Inputs to the Prefrontal Cortex. J. Neurosci. 36, 9391–9406 (2016).

207. Manoocheri, K. & Carter, A. G. Rostral and caudal basolateral amygdala engage distinct circuits in the prelimbic and infralimbic prefrontal cortex. (2022) doi:10.7554/eLife.82688.

208. Suh, J. et al. Projection-specific roles of basolateral amygdala Thy1 neurons in alcohol-induced place preference. Molecular Psychiatry 1–11 (2025).

209. Isogai, Y. et al. Multisensory Logic of Infant-Directed Aggression by Males. Cell 175, (2018).

210. de la Zerda, S. H. et al. Social recognition in laboratory mice requires integration of behaviorally-induced somatosensory, auditory and olfactory cues. Psychoneuroendocrinology 143, (2022).

211. Adolphs, R. & Spezio, M. Role of the amygdala in processing visual social stimuli. Progress in brain research 156, (2006).

212. Jabarin, R., Netser, S. & Wagner, S. Beyond the three-chamber test: toward a multimodal and objective assessment of social behavior in rodents. Molecular Autism 13, 1–29 (2022).

213. Nowlan, A. C. et al. Multisensory integration of social signals by a pathway from the basal amygdala to the auditory cortex in maternal mice. Current biology : CB 35, (2025).

214. Lopes, G. et al. Bonsai: an event-based framework for processing and controlling data streams. Front Neuroinform 9, 7 (2015).

215. Reference Atlas :: Allen Brain Atlas: Mouse Brain. https://mouse.brain-map.org/static/atlas.

216. Pnevmatikakis, E. A. & Giovannucci, A. NoRMCorre: An online algorithm for piecewise rigid motion correction of calcium imaging data. Journal of neuroscience methods 291, (2017).

217. Zhou, P. et al. Efficient and accurate extraction of in vivo calcium signals from microendoscopic video data. Elife 7, (2018).

218. Cameron, C. M., Murugan, M., Choi, J. Y., Engel, E. A. & Witten, I. B. Increased Cocaine Motivation Is Associated with Degraded Spatial and Temporal Representations in IL-NAc Neurons. Neuron 103, 80–91.e7 (2019).

219. UpSet: Visualization of Intersecting Sets. https://ieeexplore.ieee.org/document/6876017.

220. Williamson, R. C., Doiron, B., Smith, M. A. & Yu, B. M. Bridging large-scale neuronal recordings and large-scale network models using dimensionality reduction. Curr Opin Neurobiol 55, 40–47 (2019).

221. Lui, J. H. et al. Differential encoding in prefrontal cortex projection neuron classes across cognitive tasks. Cell 184, 489–506.e26 (2021).

